# Molecular Determinants for Differential Activation of the *Vibrio parahaemolyticus* Bile Acid Receptor

**DOI:** 10.1101/2022.09.02.506320

**Authors:** Angela J. Zou, Lisa Kinch, Suneeta Chimalapati, Nalleli Garcia Rodriguez, Diana R. Tomchick, Kim Orth

## Abstract

Bile acids are important for digestion of food and for antimicrobial activity. Pathogenic *Vibrio parahaemolyticus* senses bile acids via the co-component signal transduction system receptor VtrA/VtrC, an obligate membrane heterodimer. Intestinal bile acids bind to the periplasmic domain of the VtrA/VtrC complex, activating a DNA-binding domain in VtrA that induces expression of another membrane protein, VtrB. VtrB induces expression of the pathogenic Type III Secretion System 2. The bile acid taurodeoxycholate (TDC) activates VtrA/VtrC-induced *VtrB* expression, while others such as chenodeoxycholate (CDC) do not. This study demonstrates both CDC and TDC bind to the VtrA/VtrC periplasmic heterodimer using isothermal titration calorimetry (ITC). The crystal structure of the VtrA/VtrC heterodimer bound to CDC revealed it binds in the same hydrophobic pocket as TDC, but differently. Mutation of the binding pocket caused a decrease in bile acid binding affinity with exception of the S123A mutant, which bound with a similar affinity as the wild-type protein. The S123A mutant decreased TDC-induced T3SS2 activation, providing a molecular explanation for the selective activation of the T3SS2 by bile acids.

## Introduction

*Vibrio parahaemolyticus* is a gram-negative pathogen that is a leading cause of gastroenteritis from the consumption of contaminated raw seafood around the world (de Souza Santos, Salomon et al., 2015). Many cellular factors contribute to the virulence of *V. parahaemolyticus*, including two thermostable hemolysins and two Type III Secretion Systems (T3SSs), known as T3SS1 and T3SS2 (de Souza Santos et al., 2015). The T3SSs are large needle-like protein complexes that transport bacterial effector proteins, known as Vops, from the bacterial cytoplasm into the host cell cytoplasm to promote infection (de Souza Santos et al., 2015). The T3SS2 is found only in clinical isolates, and its presence coincides with gastroenteritis as well as with bacterial invasion and intracellular replication in the host gut epithelium (de Souza Santos & Orth, 2014, de Souza Santos et al., 2015, Zhang, Krachler et al., 2012). T3SS2 expression is regulated by VtrA/VtrC, a member of a newly described superfamily of bacterial “co-component” signaling systems (Kinch, Cong et al., 2022). VtrA/VtrC respond to bile acids in the host intestinal tract and induce expression of VtrB, another inner membrane transcription factor. VtrB then induces expression of the T3SS2 (Gotoh, Kodama et al., 2010, Kodama, Gotoh et al., 2010).

The newly described VtrA/VtrC-like superfamily of co-component signaling systems in enteric pathogens include members like ToxR/ToxS and TcpP/TcpH in *Vibrio cholerae* and PsaE/PsaF in *Yersinia pestis* (Kinch et al., 2022). Members of this family share several characteristics where the components of these signaling systems adopt similar domain organizations. The transcription factor component (VtrA) includes an N-terminal helix-turn-helix (HTH) DNA binding domain (DBD), a single transmembrane helix, and a C-terminal periplasmic domain, while the sensor component (VtrC) includes a transmembrane helix followed by a lipocalin-like periplasmic domain. The co-component-encoding genes share a similar operon arrangement, and their gene products form, or are predicted to form, an obligate heterodimer via interaction of the periplasmic domains (Kinch et al., 2022, Li, Rivera-Cancel et al., 2016). In the case of VtrA/VtrC, VtrA functions as a membrane-bound transcription factor, while VtrC in complex with VtrA senses bile. When VtrA/VtrC bind bile acids in the periplasm, the VtrA DBD is activated in the cytoplasm (Li et al., 2016, Okada, Matsuda et al., 2017). These characteristics distinguish co-component membrane signaling systems from a subset of membrane tethered one-component systems exemplified by the *E. coli* pH sensor CadC (Kinch et al., 2022, Schlundt, Buchner et al., 2017), as well as from two-component systems requiring an intracellular histidine kinase for signaling (Jacob-Dubuisson, Mechaly et al., 2018).

Prototypic of co-component signaling systems, the open reading frames of VtrA and VtrC overlap each other within the same operon and the two encoded proteins form an obligate heterodimer via interactions between the periplasmic domains (Li et al., 2016). Li et al. revealed this interaction using X-ray crystallography, where the apo structure of the VtrC periplasmic domain forms a β-barrel fold that incorporates a β-strand from the VtrA periplasmic domain.

They demonstrated the bile acid TDC binds to the VtrA/VtrC periplasmic domains using isothermal titration calorimetry (ITC) (Li et al., 2016), and determined the structure of the VtrA/VtrC periplasmic domain heterodimer in the TDC-bound state. In the apo structure, a mobile loop and short helix (Fα) from VtrC covers one opening of the β-barrel. This loop is displaced by TDC, which binds within a hydrophobic pocket of the VtrC β-barrel. TDC binding orders the VtrC mobile loop and extends the helix Fα, which forms a hydrogen bond to the 12α-hydroxyl (OH) of TDC.

*V. parahaemolyticus* VtrA/VtrC turns on T3SS2-mediated virulence by sensing the presence of host intestinal bile acids (Gotoh et al., 2010, Li et al., 2016). In humans, the primary bile acids cholate (CA) and chenodeoxycholate (CDC) are synthesized from cholesterol in the liver (Begley, Gahan et al., 2005, Hofmann, 1999). Prior to secretion from the liver into the intestines, the primary bile acids are conjugated to glycine or taurine producing conjugated secondary bile acids, such as taurodeoxycholate (TDC). In the distal intestines, bile salt hydrolases (BSH) from the microbiome deconjugate bile acids to regenerate free bile acids and glycine or taurine (Begley et al., 2005). Subsequently, bacterial dehydratases can remove the 7α-OH from the steroid nucleus of bile acids (Hofmann, 1999). These modifications produce the secondary bile acids deoxycholate (DC) and lithocolate. Both primary and secondary bile acids are reabsorbed into the liver and undergo re-conjugation to glycine or taurine prior to secretion back into the intestines (Begley et al., 2005, Hofmann, 1999).

Gotoh et al. previously demonstrated certain types of bile acid induce *vtrB* expression, but others do not (Gotoh et al., 2010). They grouped the various tested bile acids into three categories based on the induced level of *vtrB* transcription—high inducers, intermediate inducers, and non-inducers. Glycine-and taurine-conjugates of the secondary bile acid DC, such TDC, induced to high levels. The intermediate inducers consisted of conjugated primary bile acids and un-conjugated DC, and the primary bile acids CA and CDC did not induce. The molecular mechanism behind the variable activation of *VtrB* expression by bile acids is unknown.

Here, we use ITC to show the bile acid CDC binds to the VtrA/VtrC periplasmic complex and present the CDC-bound crystal structure. The structure shows CDC binds in the same hydrophobic pocket of the VtrA/VtrC complex as the TCD binds, and the two different bile acids compete to induce or suppress *vtrB* expression. However, TDC and CDC make different sets of interactions with the VtrA/VtrC barrel. Many mutations of residues within the bile acid binding pocket disrupt binding of TDC and CDC to VtrA/VtrC, as well as *vtrB* expression activated by TDC. However, one mutant in VtrC, S123A, did not disrupt binding but altered signaling, and the H50A mutant had marginal effects on binding, yet muted signaling. These results provide a molecular explanation for the opposite effects TDC and CDC have on inducing *vtrB* expression and the pathogenic T3SS2.

## Results

### CDC binds to VtrA/VtrC and competes with TDC

The chemical structures of the bile acids TDC and CDC share a common steroid nucleus containing three six-membered rings fused to a fourth five-membered ring (**Figure 1A)**. Attached to the steroid nucleus are a hydroxyl group at position 3α and a five carbon side chain attached to the five-membered ring. The structures of TDC and CDC differ in the position of an additional hydroxyl group attached to the steroid nucleus. TDC has a second hydroxyl group at position 12α, while CDC has a second hydroxyl group at position 7α. The carbon side chain of CDC ends with a carboxylic acid. Meanwhile, the carbon side chain of TDC is conjugated to a taurine residue via N-acyl amidation, resulting in a longer side chain (Begley et al., 2005). Due to the similarities in the structures of TDC and CDC, we reasoned CDC can bind the same hydrophobic pocket in VtrA/VtrC as bound by TDC. However, differences in the position of the hydroxyl groups and the length of the side chains in TDC and CDC may lead to different interactions with VtrA/VtrC, and thus opposing outcomes in *vtrB* transcription initiation.

**Figure 1.**
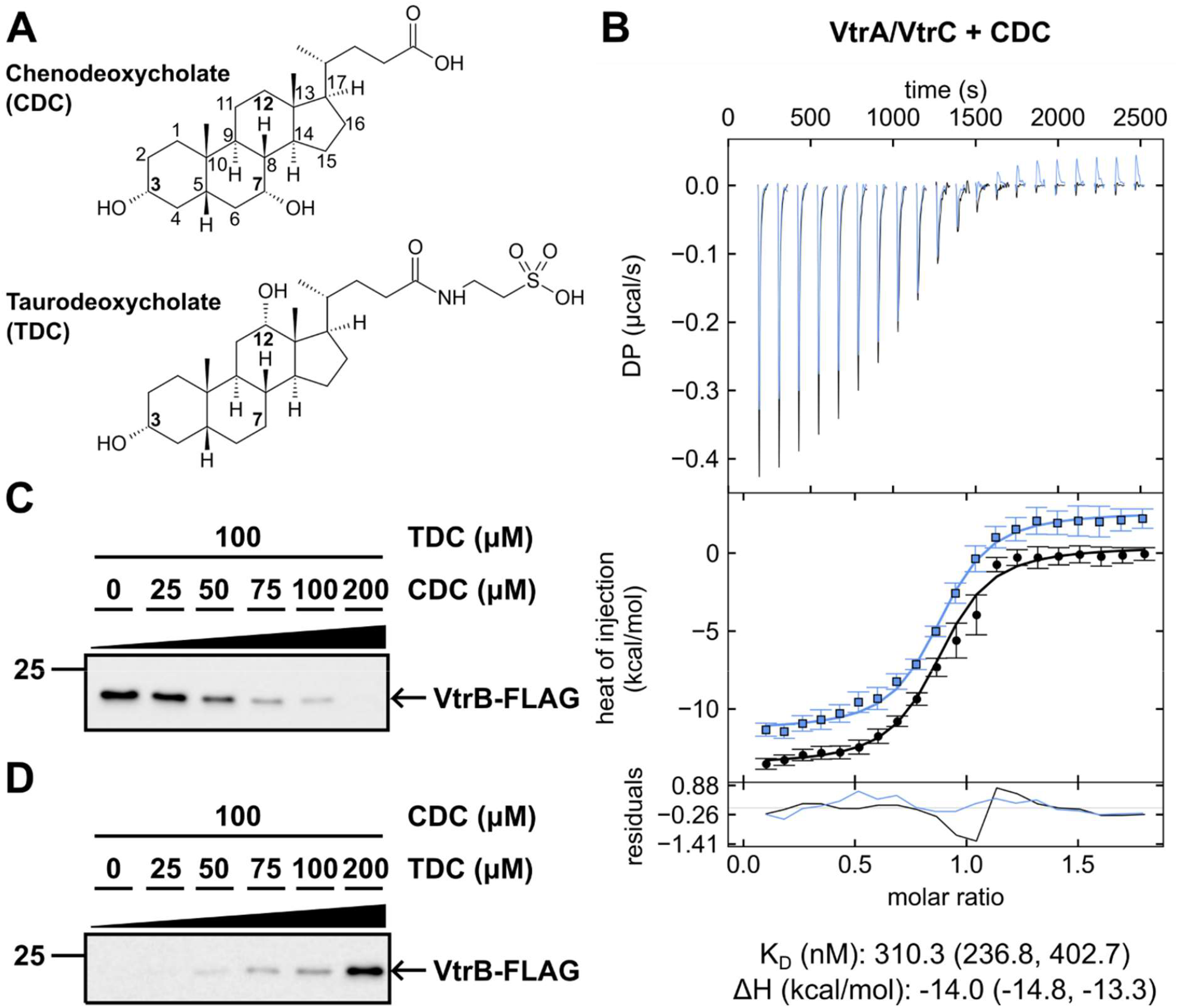
CDC binds to VtrA/VtrC and competes with TDC. **(A)** Chemical structures of CDC (Top) and TDC (Bottom). Numbers denote positions of carbon atoms in the steroid nucleus. Bold numbers indicate carbon atoms that have an attached hydroxyl group in TDC and/or CDC. Images were created in ChemSketch. **(B)** ITC thermograms for CDC binding to the VtrA/VtrC periplasmic domain complex. Thermodynamic parameters were determined by global fitting of duplicate isotherms (presented in black and blue). The dissociation constant (K_D_) and enthalpy (ΔH) values are reported with 1σ error intervals in parenthesis. **(C)** Western blot analysis of CDC/TDC competition assay. *V. parahaemolyticus* was grown in media supplemented with 100 µM TDC and varying concentrations of CDC from 0 µM to 200 µM. Anti-FLAG antibody (Sigma) was used to detect FLAG-VtrB. **(D)** Western blot analysis of TDC/CDC competition assay. Experiment was performed as in (C), except *V. parahaemolyticus* was grown in media supplemented with 100 µM CDC and varying concentrations of TDC from 0 µM to 200 µM. Results in (C) and (D) representative of three independent experiments.

We tested whether CDC binds to the VtrA/VtrC periplasmic domain complex using ITC. Like TDC (Li et al., 2016), negative power deflections were seen throughout titration of CDC into the VtrA/VtrC solution, indicating CDC binds to the complex in an exothermic manner (**Figure 1B**). The K_D_ of the CDC and VtrA/VtrC interaction was 310.3 nM with a molar ratio of approximately 1:1 (n=0.85) (**Figure 1B, Table 1**). The previously reported K_D_ of the TDC and VtrA/VtrC interaction was 315.4 nM (Li et al., 2016). When we repeated the ITC experiment with TDC and VtrA/VtrC, we obtained a lower K_D_ of 129.4 nM (**Table 1, Figure S3**). In this study, CDC bound to VtrA/VtrC with a weaker affinity compared to TDC.

**Table 1.**
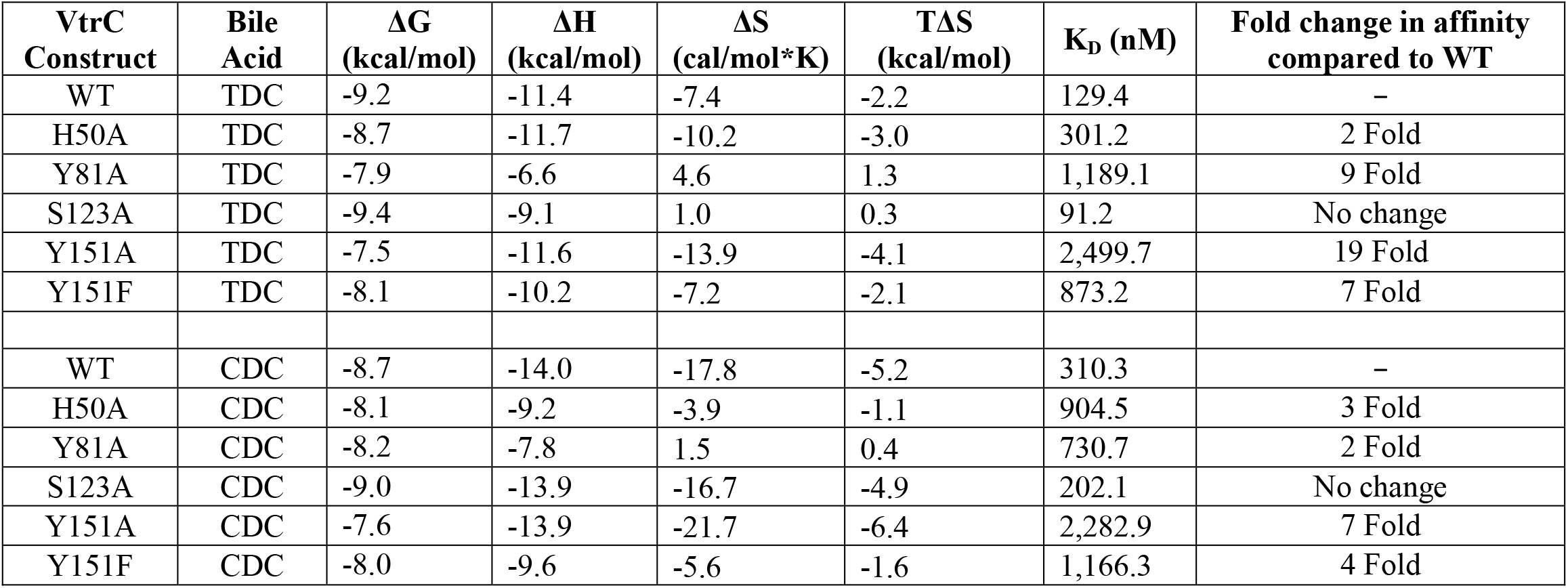
Thermodynamic parameters of TDC and CDC binding with various VtrA/VtrC constructs determined by ITC.

Gotoh and colleagues (Gotoh et al., 2010) previously demonstrated that TDC and CDC have disparate effects on the activation of VtrB. Since we observed that both CDC and TDC bind to VtrA/VtrC, we tested whether the two bile acids can compete with each other to activate *vtrB* expression. We used an attenuated *V. parahaemolyticus* clinical RimD2210633 derivative strain POR1 that is deleted for hemolysins (*ΔtdhAS*) and contains a 3XFLAG tag inserted at the 3’ end of the native *vtrB* open reading frame. We grew this strain in media supplemented with 100 µM TDC and varying concentrations of CDC from 0 µM to 200 µM. Western blot analysis of VtrB-FLAG expression revealed VtrB is expressed when treated with 100 µM TDC alone. However, VtrB expression gradually decreased with the addition of increasing amounts of CDC (**Figure 1C**). In the reverse experiment, *V. parahaemolyticus* was grown in media supplemented with 100 µM CDC and varying concentrations of TDC from 0 µM to 200 µM. VtrB was not expressed when treated with 100 µM CDC alone. VtrB expression gradually increased with the addition of increasing amounts of TDC (**Figure 1D**). These results indicate TDC and CDC compete to induce or suppress VtrB expression, respectively.

### CDC binds to VtrA/VtrC in the same hydrophobic pocket as TDC

The competitive effects of TDC and CDC on VtrB expression suggest these bile acids bind to the same region of VtrA/VtrC. We purified the VtrA/VtrC periplasmic domain complex in the presence of CDC, crystallized this complex in space group C2, and obtained the X-ray structure. The crystal structure was solved using molecular replacement with the TDC-bound structure (5KEW, space group P2_1_2_1_2_1_) as a search model and refined to a resolution of 2.08 Å. The CDC-bound VtrA/VtrC asymmetric unit contained four VtrA/VtrC heterodimers, each bound with one CDC molecule inside the β-barrel (**Figure 2A, Figure S1**).

**Figure 2.**
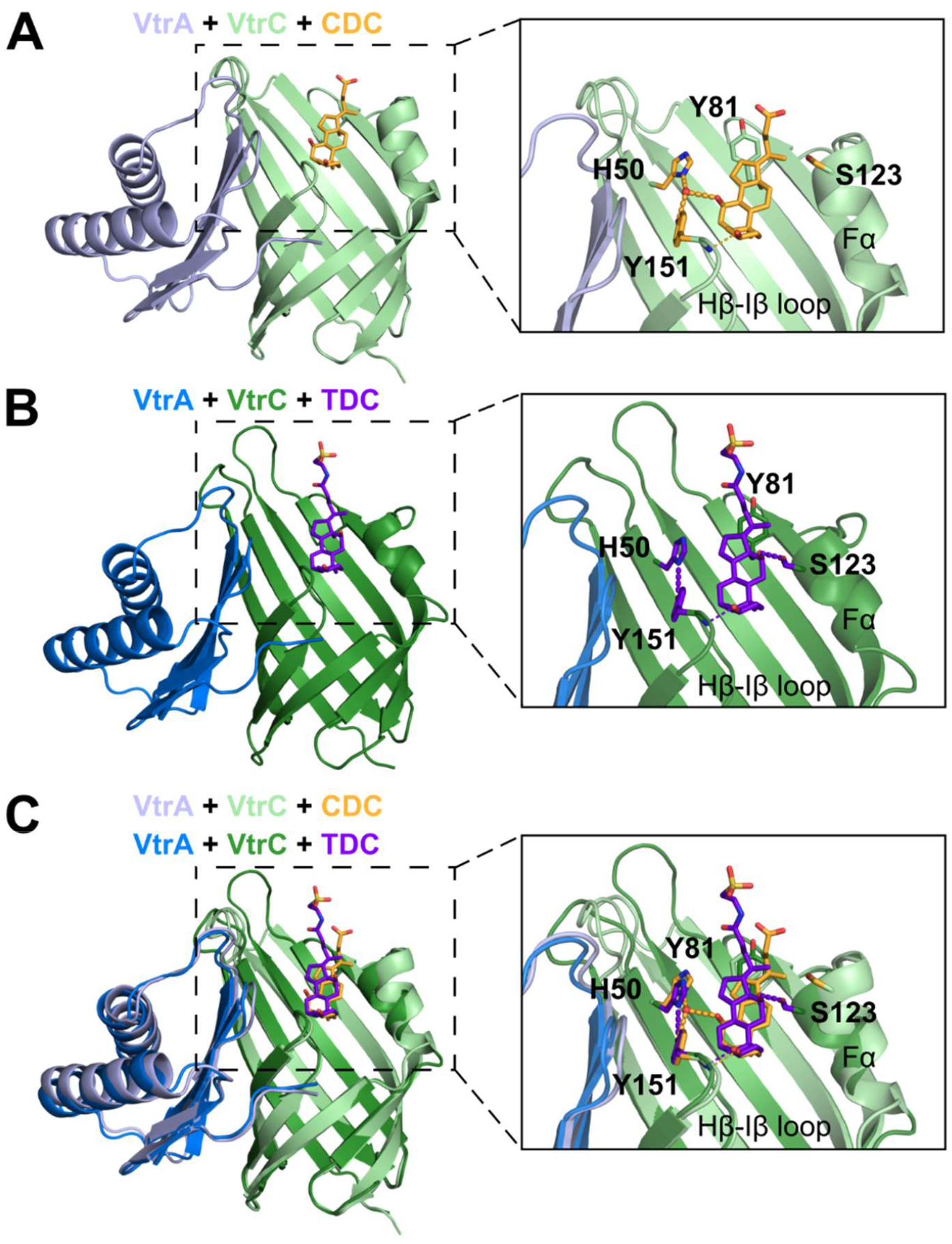
The bile acids CDC and TDC make different sets of interactions with the VtrC periplasmic domain. (**A**) Structure of the periplasmic domain complex formed by VtrA (light blue) and VtrC (light green) bound to CDC (orange). Detailed view of the CDC binding site shown in the box to the right with residues interacting with CDC and TDC shown as sticks and a coordinated water molecule represented as a red sphere. Hydrogen bonds between protein, CDC, and water are represented as orange dashed lines. (**B**) Structure of the VtrA (dark blue) and VtrC (dark green) complex bound to TDC (purple). Detailed view of the TDC binding site shown in the box to the right with the same residues as in (**A**) shown as sticks. Hydrogen bonds between protein and TDC are represented as purple dashed lines. (**C**) Overlay of the images in (**A**) and (**B**) showing differences in the structures of VtrA/VtrC, positions of interacting residues, and hydrogen bonds between the protein and CDC/TDC. Images in (A, B, and C) were generated from the structures of the CDC- and TDC-bound heterodimers with the lowest mean temperature factors in each crystal.

Despite their common steroid nucleus, CDC is modified with a 7α-OH that is absent in TDC (**Figure 1A**). This unique CDC hydroxyl group forms a hydrogen bond with a water molecule that is also coordinated by VtrC H50 ND1 and Y151-OH (**Figure 2A**). The backbone amide of VtrC Y151 forms a hydrogen bond with the 3α-OH of CDC, and VtrC Y81 makes van der Waals contacts with CDC. Electron density for CDC and the surrounding binding pocket residues was more defined in two of the four VtrA/VtrC heterodimers (**Figure S1**). The models for the heterodimers with lower atomic displacement parameters and more defined density included a water molecule in the binding pocket, while the other heterodimers did not. However, the distance between H50 ND1 and Y151-OH was very similar in all four heterodimers, ranging from 4.7Å – 5.0 Å. The CDC binding site overlaps with the previously determined TDC site, with the taurine conjugate facing outside the barrel (**Figure 2B**), The steroid backbone of each bile acid overlaps in a superposition of the two structures, with the conjugated side chain position exhibiting the largest deviation (**Figure 2C**). The relative position of VtrA is similar in both structures.

Alignment of all four CDC-bound heterodimers in the asymmetric unit showed a difference in the position of the short VtrC helix Fα in one heterodimer compared with the other heterodimers (**Figures 3A, S2A**). Fα was located further from the CDC binding site in one heterodimer (chains A/B). This helix forms direct contacts with a symmetry mate in the crystal lattice. In the other three heterodimers, Fα does not contact symmetry mates. Meanwhile, on the other side of the VtrA/VtrC β-barrel, the VtrC Hβ-Iβ loop forms part of the interface with VtrA (**Figure 2A**). Within this loop is VtrC Y151, which forms a hydrogen bond with CDC 3α-OH. Residue Y151 is aligned closely near the VtrA interface in all four CDC-bound heterodimers (**Figure S2A**). Also near the VtrA/VtrC interface is a flexible loop containing VtrC residues 43-48 (**Figure S2A**). This loop adopts relatively similar positions in all four CDC-bound heterodimers. However, this loop does not make polar contacts with VtrA **(Figure S2C)**. The root mean square deviation (RMSD) of VtrA among the four heterodimers ranged from 0.3Å – 0.8Å, while the RMSD of VtrC ranged from 0.5Å – 0.8Å (**Figure S2A, right panel**). VtrA and VtrC in chains A and B had the largest RMSDs compared with the other three heterodimers, ranging from 0.7 – 0.8 Å for each.

**Figure 3.**
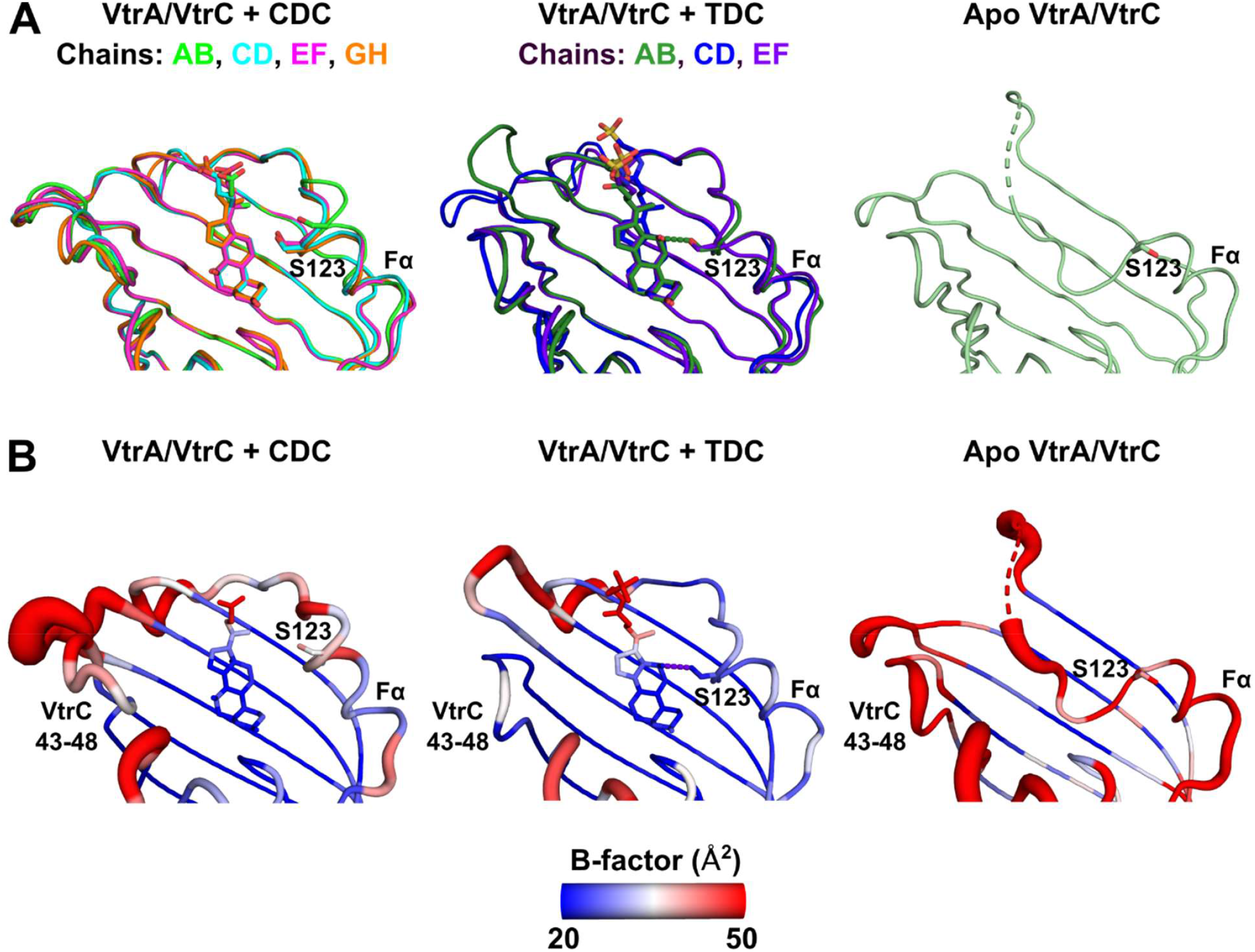
Hydrogen bond between TDC and S123 constrains Fα helix. **(A)** Superposition of all four heterodimers in the asymmetric unit of the CDC-bound crystal (left) and all three heterodimers in the asymmetric unit of the TDC-bound crystal (middle). Apo heterodimer (left). Structures of VtrA/VtrC represented by ribbons. CDC, TDC, and S123 are shown as sticks. S123 and the Fα helix are labeled. Hydrogen bond between TDC and S123 in each heterodimer shown as dashed lines. **(B)** Representation of the atomic displacement factors (i.e., “B-factors”) showing the Fα helix has higher B-factors in the CDC-bound and apo structures than in the TDC-bound structure.

### Binding pocket changes are observed with CDC or TDC and VtrA/VtrC

The previously determined TDC-bound structure revealed a key interaction with VtrC S123 that ordered part of the helix Fα in the presence of ligand (Li et al., 2016). In the TDC-bound structure, the 12α-OH of TDC makes a hydrogen bond with the VtrC S123-OH (**Figure 2B**). VtrC H50-ND1 and Y151-OH form a direct hydrogen bond, instead of coordinating a water molecule as is observed in the CDC-bound structure (**Figure 2**). The backbone amide of Y151 forms a hydrogen bond with TDC 3α-OH, similar to that observed in CDC. Y81 forms van der Waals contacts with the carbon sidechain from TDC.

A superposition of the TDC- and CDC-bound heterodimers revealed differences in ligand interactions (**Figure 2C**). Specifically, the coordinated water molecule between H50, Y151, and the 7α-OH of CDC increases the distance between the H50 and Y151 sidechains, as compared to the TDC-bound structure, where they form a hydrogen bond. The key Fα S123 sidechain makes a hydrogen bond with 12α-OH of TDC, which is lacking in CDC (**Figure 2C**). Finally, the Y81-OH group shifts between 0.8 Å and 2.3Å in the CDC bound structure so that the aromatic sidechain forms van der Waals contacts with both the five-membered ring and the carbon sidechain from CDC (**Figure 2C**).

The residues from the Fα helix are variable in a superposition of the four heterodimers in the asymmetric unit of the CDC-bound crystal (**Figure 3A, S2A**), and they and have relatively higher atomic displacement parameters (i.e., “B-factors”) than in the TDC-bound crystal (**Figure 3B**). The Fα helix residue S123 shifts away from the CDC binding site in one heterodimer compared to the other three due to crystal packing (**Figures 3A, S2A**). In the TDC-bound crystal, S123 forms a hydrogen bond with TDC in two of the three heterodimers in the asymmetric unit (**Figure 3A**). In the other heterodimer (chains C/D), part of the Fα helix containing S123 and an unstructured loop are not modeled due to poor electron density (Li et al., 2016). While the modeled Fα helix in this chain C/D heterodimer aligns closely with that of the chain E/F heterodimer, the Fα helix of chain A/B differs as a result of an interaction with the neighboring VtrA (chain C) in the asymmetric unit. Despite these conformational differences, the TDC bound B-factors are lower than in the CDC-bound heterodimers (**Figure 3B**). These superpositions suggest the hydrogen bond between TDC and S123 constrains the Fα helix near the binding pocket. Meanwhile, the lack of a hydrogen bond between CDC and S123 allows mobility in the Fα helix and alternate conformations dictated by the crystal contacts.

The positions of two loops surrounding the bile acid binding site of the VtrA/VtrC β-barrel near Y81 (VtrC residues 70-79) and near the interface with VtrA (VtrC residues 43-48) also differed in the CDC- and TDC- bound structures (**Figure 2C, Figure S1**). These loops adopt different conformations due to interactions with other heterodimers in the asymetric unit as well as interactions from crystal packing. The interface proximal loop containing VtrC residues 43-48 adopts distinct positions in the CDC- and TDC-bound structures (**Figure S2C**). In the CDC-bound structure, this loop maintains a similar orientation in all four heterodimers. None of the sidechain residues from the CDC-bound loop make polar contacts with VtrA. In the TDC-bound structure, this loop rotates in all three heterodimers such that VtrC D45 moves towards a flexible loop in VtrA (residues 234-239). In one of the TDC-bound heterodimers, VtrC D45 forms a hydrogen bond with the backbone amide of VtrA E235 (**Figure S2C**). Additionally, the loop containing VtrC residues 43-48 has higher B-factors in the CDC-bound structure compared to in the TDC-bound structure (**Figure 3B**), indicating this loop is more mobile in the CDC-bound complex. Given the proximity to VtrA and the consistently alternate conformations displayed by all CDC-bound and TDC-bound heterodimers, the interaction between this loop and VtrA may be important for TDC induced activation of *vtrB* expression.

### Pocket residues in VtrC dictate its binding to bile acids

To test the contribution of residues in the bile acid binding pocket to TDC and CDC binding, we made mutant constructs of the VtrA/VtrC periplasmic domains so that their binding to bile acids could be compared to wild-type (WT) binding (**Figure 1B**). The mutants contain single amino acid substitutions of residues H50, Y81, S123, and Y151 with alanine or phenylalanine (**Table 1**). We then tested the binding of TDC and CDC to these VtrA/VtrC mutants. Most of the VtrA/VtrC mutants bound both TDC and CDC less favorably compared WT, as shown by 2 fold (H50A) to 19 fold (Y151A) increases in K_D_ for TDC and 2 fold (Y81A) to 7 fold (Y151A) increases in K_D_ for CDC (**Table 1**). The exception to this trend was the S123A mutant, which had a similar K_D_ for both TDC and CDC compared to WT.

Using the ITC technique to measure bile acid binding is advantageous in that it provides thermodynamic parameters of entropy (ΔS) and enthalpy (ΔH) in addition to binding constants. Considering the values of these parameters in light of the apo and bile acid bound structures can provide insight into how binding is mediated by interactions (from enthalpy, ΔH) and disorder (from entropy, TΔS). The enthalpy change of WT binding to TDC (ΔH = −11.4 kcal/mol) is less favorable than its binding to CDC (ΔH = −14.0 kcal/mol), while the entropic contribution of binding TDC (TΔS = −2.2 kcal/mol) is more favorable than binding CDC (TΔS = −5.2 kcal/mol) (**Figure S4**). Consistent with these entropic values, a mobile loop from the apo structure (**Figure S4A, orange tube**) is ordered upon bile acid binding, and B-factors from the apo structure (**Figure 3B**) are generally higher than those in the CDC-bound and TDC-bound structures. The more restricted freedom resulting from the WT VtrA/VtrC binding to CDC can not be explained by the structure B-factors, suggesting the difference arises from additional ordering of solvent in the CDC bound state. Accordingly, the CDC binding specificity results from an ordered water interaction with the 7α-OH (**Figure S4C**). Additional contributions to the entropic difference could include the increased flexability of the bound TDC with its carbon tail sidechain conjugated to taurine with respect to the shorter unconjugated carbon tail sidechain for CDC, or from differential higher order protein interactions in the CDC and TDC bound states.

In addition to the thermodynamic differences in WT bile acid binding, the VtrC mutations had varying effects on the enthalpic and entropic contributions to binding. The Y81A mutation decreased the enthalpy change for both TDC and CDC binding when compared to WT, as well as favorably altered the entropic contributions to positive values (**Table 1**). The difference in ΔH resulting from the Y81A mutation could result from a decrease in van der Waals contacts between either of the bile acids and the smaller alanine side chain mutation compared to that with the WT tyrosine 81 side chain (**Figure 2**). The increased entropy of the Y81A mutation suggests the Y81 sidechain promotes the stability of the bound state of the protein when compared to the free state, possibly through bile acid-induced organization of the mobile loop (**Table 1**). The entropy difference observed for TDC binding with respect to CDC binding to the Y81A mutant (**Table 1**) may reflect the observed ordering of Fα helix by S123 upon binding TDC (**Figure 2B, Figure 3B**).

### A conformational switch in the binding pocket accomodates CDC

Mutation of H50 and Y151, whose side chains form a hydrogen bond in the WT TDC-bound structure and line the binding pocket (**Figure 2B, Figure S4**), should theoretically decrease the enthalpy of binding to the bile acids. However, both H50A and Y151A had similar ΔH for TDC binding as the WT protein (**Table 1**). The apo structure of VtrA/VtrC experiences a conformational change, where an extended loop from the Fα helix covers the bile acid binding site. In the apo structure, the H50/Y151 pair forms the same hydrogen bond as in the TDC bound state, but they instead interact with P121 from this loop. The hydrophobic ring of P121 is replaced by the middle six-membered ring from the TDC steroid nucleus upon binding (**Figure S4**). Thus, the similar ΔH values for the mutant and WT proteins likely reflect a similar entropic contribution to binding the Fα helix loop in the apo state as to binding the steroid nucleus in the TDC-bound state. The enthalpic energies for each state (apo and TDC-bound) in the mutant should increase to the same extent (**Figure S4**). Mutation of the H50/Y151 hydrophobic anchor for the Fα extended loop should also increase its flexibility. The elevated changes in entropy for TDC binding to the H50A and Y151A mutants support this notion.

The Y151A mutant also displayed similar ΔH as WT when binding CDC, but the ΔH value for the H50A mutant was decreased (**Table 1**). In the CDC-bound structure, the pocket changes to accomodate the 7α -OH group of CDC. The position of the H50 sidechain switches to form a hydrogen bond with an ordered water instead of the Y151 side chain OH. The ordered water establishes a hydrogen bond network between H50, Y151, and the 7α -OH from CDC (**Figure 2A, Figure S4C**). Because the conformational switch occurs only for CDC binding, the decreased ΔH of the H50A mutation with respect to WT suggests the CDC-bound state has an increased enthalpic energy compared to the TDC bound state that represents the conformational switch (**Figure S4**).

The Y151F mutation distinguishes between the ability of the sidechain to form hydrogen bonds from its aromatic contribution to hydrophobic interactions. The Y151F mutation altered the ΔH of TDC binding less than that of CDC binding compared to WT, where the contribution of enthalpy decreased (**Table 1**). This decrease would be consistent with the loss of the Y151 hydrogen bond, destabilizing the CDC bound state where the hydrogen bonding network accomodates the 7α-OH (**Figure 2A**).

### Residue S123 dictates bile acid specificity for inducing *VtrB* transcription

The S123A mutation did not change the binding affinity of VtrA/VtrC to either bile acid (**Table 1**). However, the ΔH of TDC binding decreased and TΔS shifted to a positive value, likely due to the loss of a stabilizing hydrogen bond between the TDC 12α-OH and S123 on the Fα helix (**Figure 2B, Figure S4**). Alternately, the ΔH and TΔS of CDC binding was similar to that of the WT protein because S123 does not form a hydrogen bond with CDC (**Figure 2A, Figure S4**). Because binding to bile acids does not change for the S123A mutant, we suspected the conformation of the Fα helix imposed by the S123 sidechain might dictate the specificity of the VtrB transcription factor activation. To test this hypothesis, we used a GFP reporter assay in which a *V. parahaemolyticus* POR1*ΔvtrC* strain (**see Materials and Methods**) expressed N-terminally FLAG-tagged *vtrC* (WT) or single amino acid substitutions of *vtrC* (including Q42A, H50A, Y81A, S123A, Y151A, and Y151F) from a plasmid under the control of an arabinose inducible promoter. The strains also contained a second plasmid with the *gfp* gene under the control of the *V. parahaemolyticus vtrB* promoter (Okada et al., 2017) with the 300 bp upstream region of the *vtrB* open reading frame (P_*vtrB*_*-gfp*). The strains were treated with either 100 μM TDC or 100 μM CDC, and GFP fluorescence intensity and absorbance at 600 nm (A_600_) were measured every five minutes for one hour. The GFP fluorescence readings were then normalized by A_600_ and the fluorescence readings at the one hour time point were plotted in **Figure 4**.

**Figure 4.**
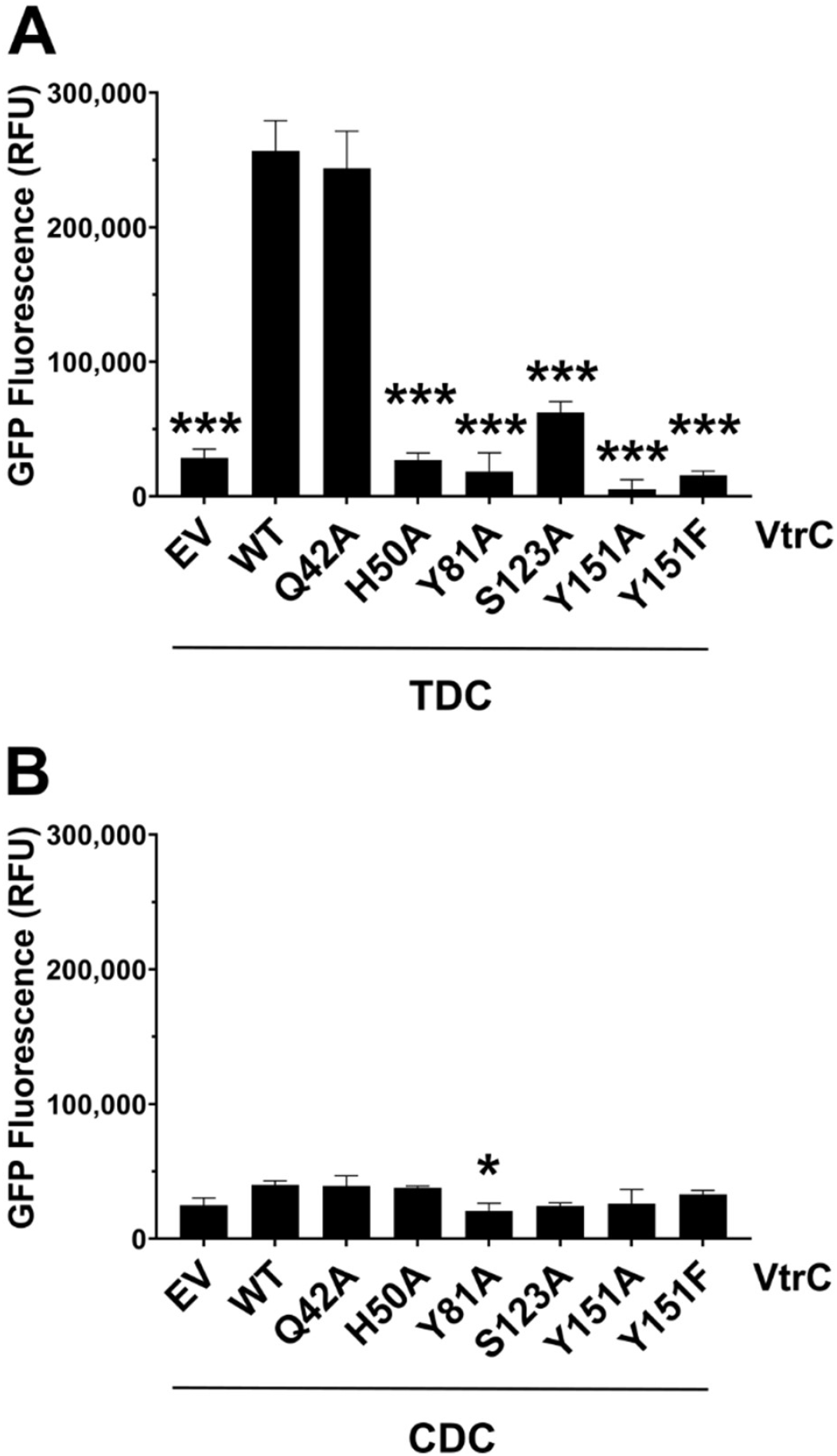
Mutation of residues involved in bile acid binding disrupt TDC-induced signaling, while CDC does not induce signaling. **(A)** GFP fluorescence from a P_*vtrB*_*-gfp* transcriptional reporter in *V. parahaemolyticus* POR1Δ*vtrC* strains expressing FLAG-VtrC (WT) and single amino acid substitution mutants upon treatment with TDC. EV, empty vector control. **(B)** GFP fluorescence of POR1Δ*vtrC* strains in (A) treated with CDC. Results are the mean of three technical replicates. Results are representative of three biological replicates. Error bars denote SD. *, p < 0.05; ***, p < 0.0001; comparisons to the WT strain determined by one-way ANOVA followed by Tukey’s multiple comparison test.

Substitution of S123 with alanine resulted in a significant four-fold decrease in GFP fluorescence in response to TDC compared to the WT strain, suggesting this residue plays a role in transcription factor activation in response to binding TDC. In the presence of CDC, the S123 mutant exhibited similar low levels of GFP fluorescence that was not significantly different than the empty vector (EV). The WT-level binding of the S123A mutant to both TDC and CDC, together with the selective activation of *VtrB* transcription in response to TDC, but not CDC, supports a role for the S123 hydrogen bond stabilization of the Fα helix in TDC-specific activation of the VtrA/VtrC transcription factor.

### Bile acid binding residues induce *VtrB* transcription

Substitution of the remaining binding residues H50, Y81, and Y151 with alanine or phenylalanine all resulted in greater than 75% decrease in GFP fluorescence in response to TDC compared to the WT strain (**Figure 4A**). Meanwhile, the Q42A mutation, located outside of the binding pocket (Li et al., 2016), had similar GFP fluorescence as the WT strain. Similar to the results for S123A, treatment of the WT and mutant strains with CDC induced GFP fluorescence at low levels that were not significantly different from an empty vector (EV) control (**Figure 2B**). These results indicate the tested binding pocket residues are all important for activation of *VtrB* transcription in response to TDC. Meanwhile, CDC does not induce transcription of *VtrB* in any of the WT or mutant strains.

### Residue H50 is important for *VtrB* transcription under acidic and neutral conditions

*V. parahaemolyticus* encounters greatly different pH environments as the bacterium travels through the human gastrointestinal (GI) tract after being consumed. The human GI tract varies from pH 1.0 – 2.0 in the stomach to pH 6.6 – 7.5 in the small intestines and pH 6.5 – 7.0 in the large intestines (Evans, Pye et al., 1988). Since histidine residues can have different protonation states under various physiologic pH conditions, we tested whether pH alters the activity of VtrA/VtrC in response to TDC.

We cultured the WT, H50A, and S123A strains and performed the GFP reporter assay in media adjusted to pH 5.5 using hydrochloric acid, pH 7.0 (no adjustment), and pH 9.0 using sodium hydroxide. For the WT strain, the level of GFP fluorescence at pH 5.5 and pH 7.0 was not significantly different, while it decreased at pH 9.0 (**Figure 5**). For the H50A strain, the GFP fluorescence was lower than that of the WT strain under each pH condition. However, the decrease in GFP fluorescence shifted to a lower pH (between pH 5.5 and pH 7.0). The fluorescence at pH 7.0 was not significantly different from the fluorescence at pH 9.0. A similar trend was seen for the S123A mutant, except the fluorescence at pH 7.0 was significantly higher than at pH 9.0. These results indicate H50 and S123 are important for activating *vtrB*transcription under acidic and neutral conditions.

**Figure 5.**
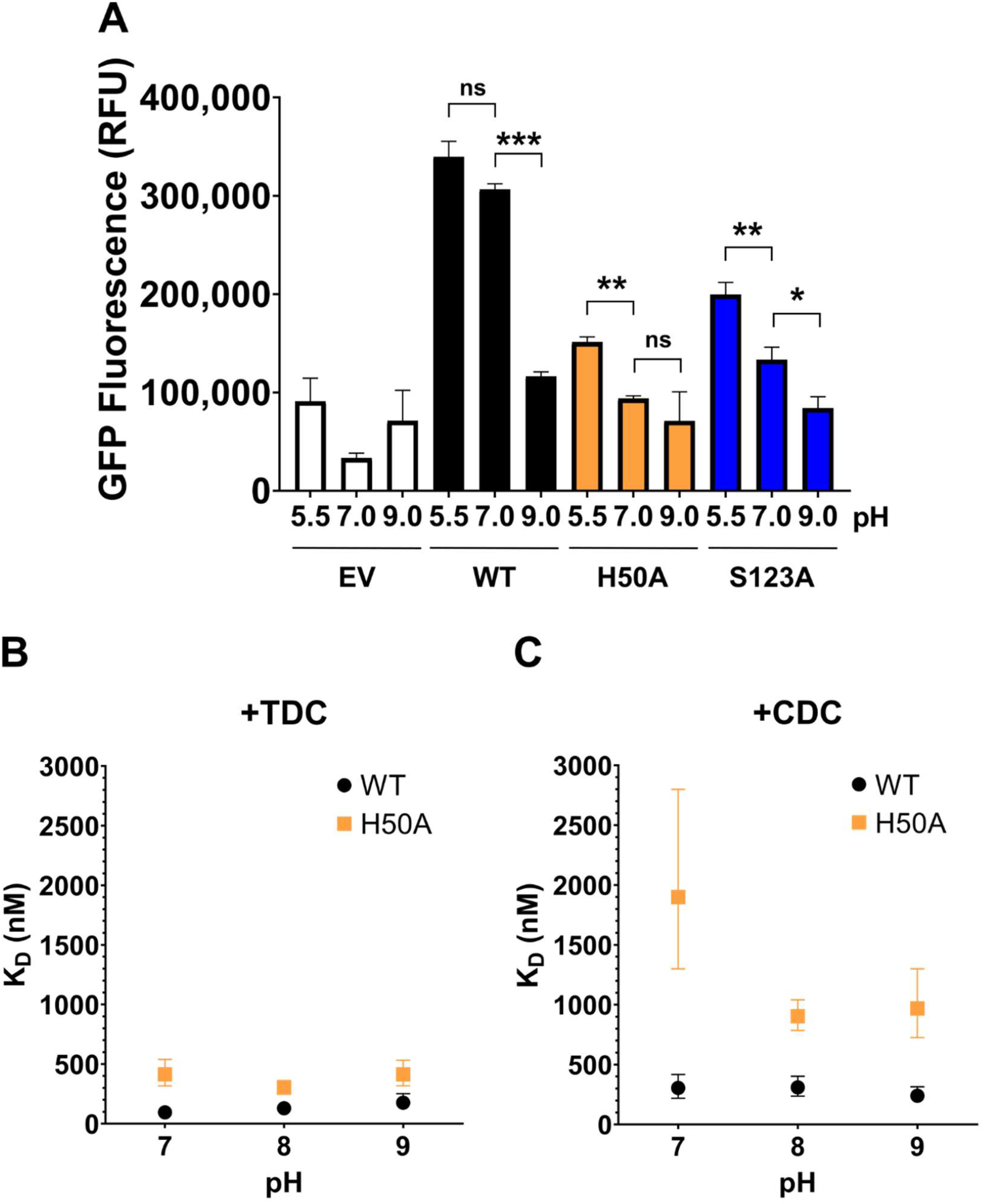
H50A and S123A mutations reduce ability of TDC to induce transcription under acidic and neutral conditions. (**A**) GFP fluorescence from a P_*vtrB*_*-gfp* transcriptional reporter in *V. parahaemolyticus ΔvtrC* strains expressing VtrC (WT) and single amino acid substitution mutants upon treatment with 100 μM TDC under different pH conditions. Results are the mean of three technical replicates. Error bars denote SD. ns, non-significant; *, p < 0.05; **, p < 0.005; ***, p < 0.0001; determined by one-way ANOVA followed by Tukey’s multiple comparison test. (**B**) K_D_ of TDC binding WT and H50A protein constructs under various pH conditions measured by ITC. Error bars denote 68.3% confidence intervals. (**C**) K_D_ of CDC binding WT and H50A protein constructs under various pH conditions measured by ITC. Error bars denote 68.3% confidence intervals.

We reasoned the different levels of *vtrB* transcription under different pH conditions may be a result of pH-dependent changes in the binding affinity of TDC and CDC. To test the effect of pH on binding affinity, we performed additional ITC experiments with the VtrA/VtrC periplasmic domains. The ITC results in **Table 1** were measured using a pH 8.0 buffer (**see Materials and Methods**). We then purified the WT and H50A constructs of the VtrA/VtrC periplasmic domains in pH 7.0 and pH 9.0 buffers. The binding affinities of TDC and CDC to each protein construct were measured using these same buffers at pH 7.0 and pH 9.0. The K_D_ measurements versus pH for TDC and CDC are plotted in **Figure 5B-C**, respectively. The WT protein bound TDC with slightly higher K_D_ with increasing pH (**Figure 5B**). However, these K_D_ values were all within the 68.3% confidence intervals (CIs) of each other and were not significantly different (**Supplemental Table S3**). Similarly, the WT protein bound CDC with similar affinities under each pH condition (**Figure 5C**). Meanwhile, the H50A construct bound TDC with similar affinities under each pH condition, with higher K_D_ values compared to that of WT with TDC (**Figure 5B**). Interestingly, the H50A construct bound CDC with a higher K_D_ around 1,900 nM at pH 7.0 compared to K_D_ values of around 900 nM under pH 8.0 and pH 9.0 conditions (**Figure 5C**). However, the lower limit of the 68.3% CI of the K_D_ at pH 7.0 was close to the upper limit of the corresponding CI at pH 9.0 (**Supplemental Table S3**). As a result, we cannot confidently say the binding affinity of H50A with CDC is significantly different at pH 7.0 compared to pH 8.0 and pH 9.0.

These ITC results indicate the bile acid binding affinities of the WT and H50A constructs do not change significantly under different pH conditions. The different levels of *VtrB* transcription under various pH conditions (**Figure 5A**) is not a result of changes in bile acid binding. Instead, residues outside of the bile acid binding site may control the differential activity of the complete VtrA/VtrC protein in response to pH. Additional studies are needed to understand the differential activation of VtrA/VtrC in response to pH.

## Discussion

Herein, we demonstrate the molecular determinants of TDC and CDC that dictate whether or not the co-component signaling system VtrA/VtrC can transcriptionally activate *VtrB* and the pathogenic T3SS2 (**Figure 7**). CDC and TDC bind to the same hydrophobic pocket in the VtrA/VtrC periplasmic domain complex, but form different interactions with binding pocket residues. Functional and binding assays of various VtrA/VtrC mutants showed the hydrogen bonds between VtrC H50 and Y151 and the TDC 12α-OH and S123 are important for activating *VtrB* transcription (**Figures 2B, 4A)**. The van der Waals interaction between TDC and Y81 is also important for indiscriminant binding of the bile acids, and its absence supresses transcription. In the case of CDC binding, the H50 sidechain switches conformation to allow hydrogen bonding to an ordered water molecule in between these two residues and the 7α-OH specific to CDC (**Figure 2A, Figure S4C**). There is no interaction between S123 and CDC, because CDC has a 7α-OH instead of a 12α-OH (**Figure 1A**). The ability of S123 to stabilize the Fα helix and extended loop when bound to TDC, but not CDC might explain the specificity between the two bile acids for activating *VtrB* transcription. Mutation of this residue does not alter binding to TDC, but instead increases the entropy (**Table 1**) and reduces the ability of VtrA/VtrC to activate transcription of VtrB (**Figure 4**). Alternately, the TΔS of the S123A mutant does not change much for CDC, which does not activate transcription. The VtrA/VtrC transcription factor might harness the lower entropy of the S123 stabilized Fα helix in the TDC bound state to dimerize for transcription activation. The structure determination and ITC binding experiments in this study are limited to the periplasmic domains of VtrA/VtrC, and pH response of binding to the periplasmic region compared to transcription activation by the full length VtrA/VtrC differs. More studies are needed to determine the molecular mechanism of the inducing activity of TDC and non-inducing activity of CDC in the context of full-length VtrA/VtrC.

The model pKa of the histidine side chain, which is the pKa value of histidine in water, is well known and measured to be 6.50 (Li, Robertson et al., 2005). However, the pKa of ionizable residues in a protein can vary due to desolvation effects from the burial of the residue in the protein and intra-protein interactions with nearby residues (Li et al., 2005). The empirical pKa predictor PROPKA 3.4.0 (Olsson, Søndergaard et al., 2011, Søndergaard, Olsson et al., 2011) suggests this to be the case for the H50 switch residue. The predicted pKa of H50 was 3.75 in the apo structure, 3.68 in the TDC-bound structure, and 5.44 in the CDC-bound structure. The apo structure was crystallized in pH 5.6 buffer, the TDC-bound structure in pH 4.6 buffer (Li et al., 2016), and the CDC-bound structure in pH 7.0 buffer (**see Methods and Materials**).

To predict the pKa shift caused by the desolvation effect, PROPKA 3.4.0 calculates how buried a residue is in the protein structure. The desolvation effect raises the pKa of buried acidic residues and lowers the pKa of buried basic residues (Olsson et al., 2011). According to PROPKA 3.4.0, VtrC H50 was 77% buried in the apo structure, 75% buried in the TDC-bound structure, and 63% buried in the CDC-bound structure. These % burial values correspond with the burial of H50 seen in the crystal structures. In the apo structure, H50 is covered by the flexible loop containing the short Fα helix (**Figure 6A**). In the TDC-bound structure, this loop is displaced, but H50 is covered by TDC binding (**Figure 6B**). In the CDC-bound structure, the position of H50 is shifted slightly away from Y151 due to the interaction with a coordinated water molecule (**Figure 2C**) and H50 is more exposed (**Figure 6C**).

**Figure 6.**
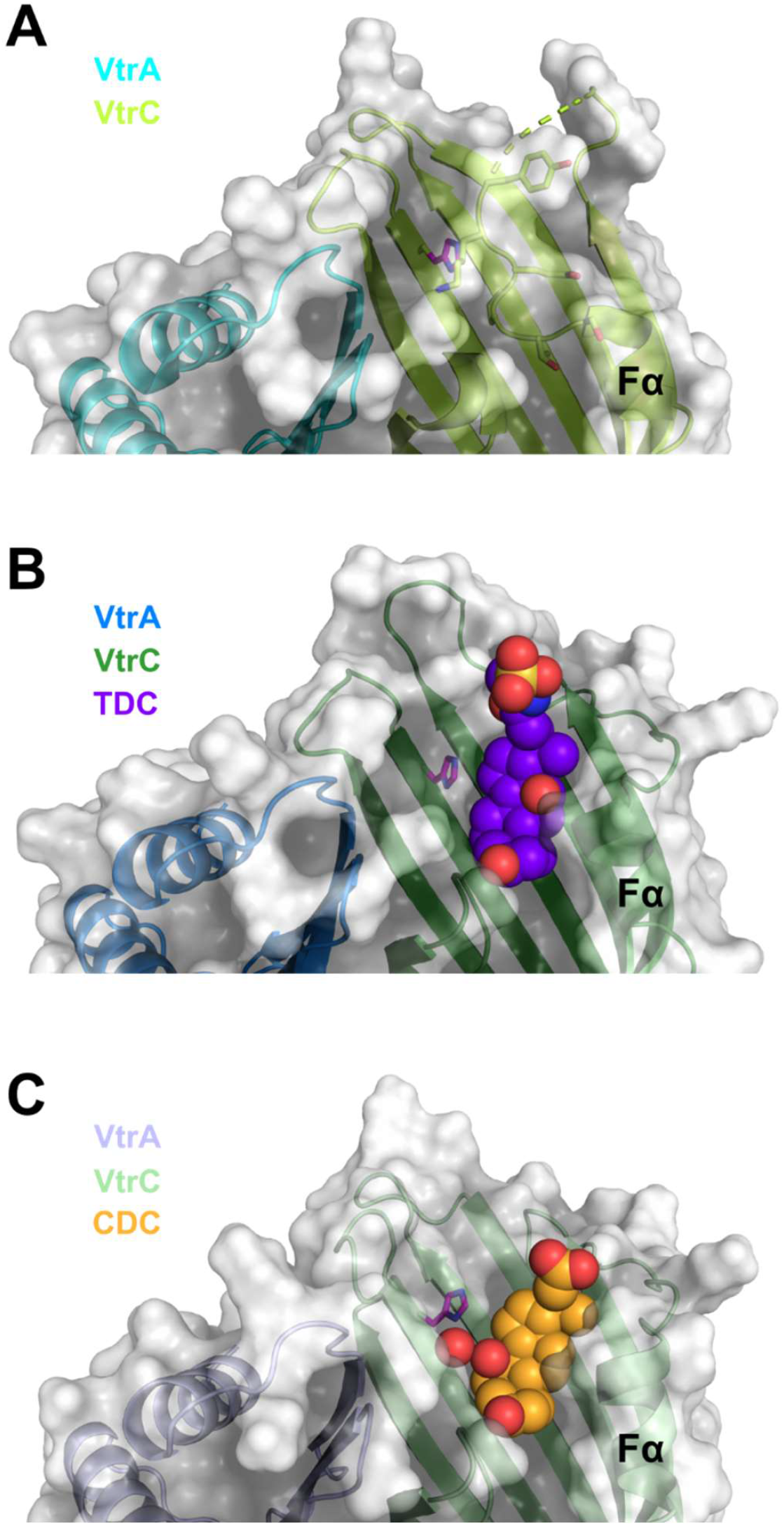
Burial of VtrC H50 in the apo, TDC- and CDC-bound structures. **(A)** Overlay of surface and ribbon models of apo VtrA/VtrC. Side chains of H50 and residues in the flexible loop covering binding pocket shown as sticks. H50 highlighted in magenta. **(B)** Overlay of surface and ribbon models of TDC-bound VtrA/VtrC. TDC shown as spheres. Side chain of H50 shown as sticks and highlighted in magenta. **(C)** Overlay of surface and ribbon models of CDC-bound VtrA/VtrC. CDC and coordinated water molecule shown as spheres. Side chain of H50 shown as sticks and highlighted in magenta.

**Figure 7.**
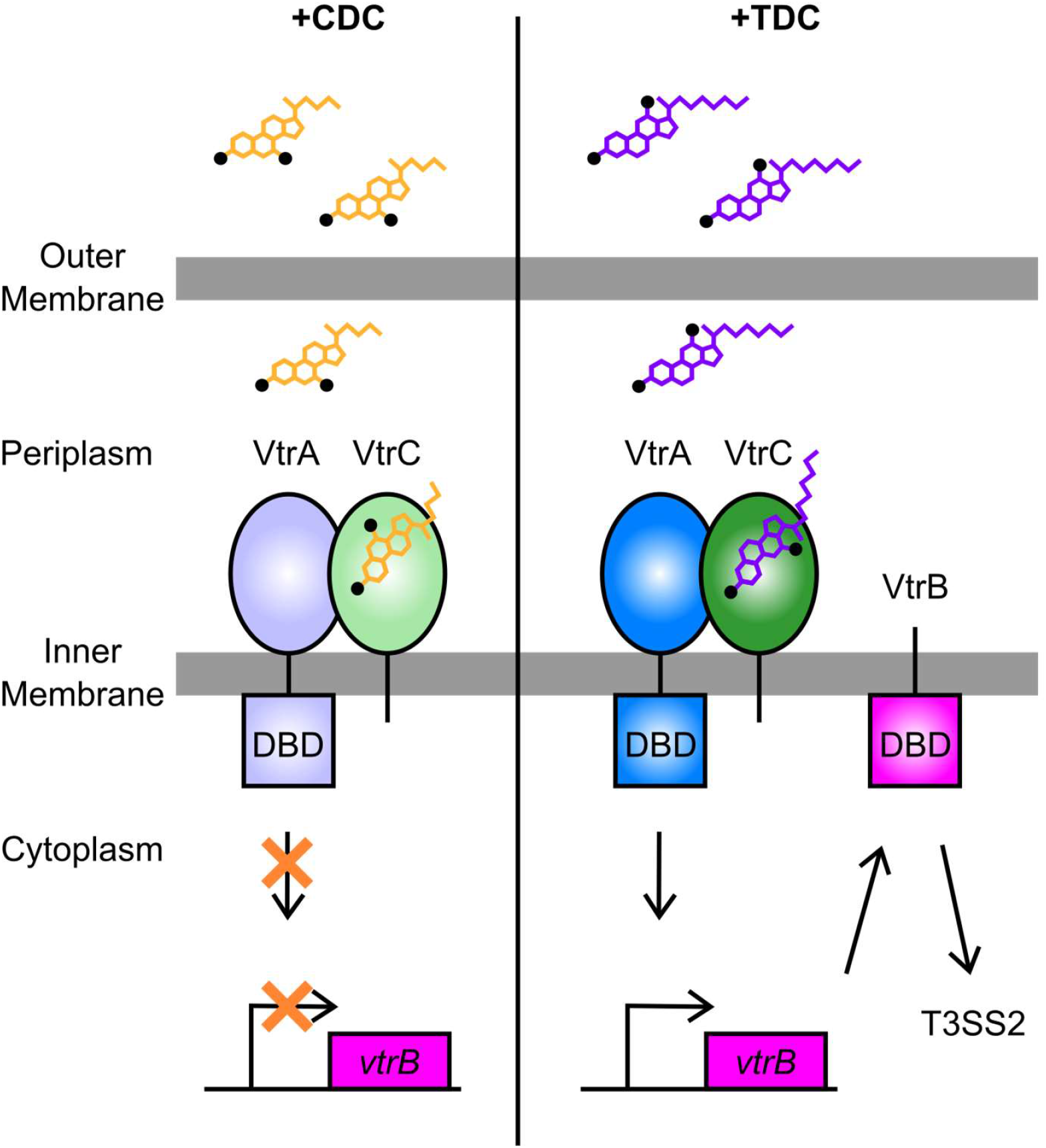
CDC and TDC binding to VtrA/VtrC produces different outcomes on *vtrB* transcription. **(Left)** When the bile acid chenodeoxycholate (CDC) binds to the periplasmic domain of VtrC, the cytoplasmic DNA binding domain (DBD) of VtrA does not induce transcription of *vtrB*. **(Right)** In contrast, when taurodeoxycholate (TDC) binds to the periplasmic domain of VtrC, the VtrA DBD induces transcription of *vtrB*, leading to the expression of another transmembrane protein containing a cytoplasmic DBD, VtrB. Subsequently, VtrB induces expression of Type III Secretion System 2 (T3SS2) encoding genes. Hydroxyl groups attached at different positions on CDC and TDC depicted as black dots.

Based on the similar predicted pKa values and percentage burial of H50 in the apo and TDC-bound structures, we speculate the H50-Y151 interaction (**Figure 2B**) primes VtrA/VtrC for activation. Upon displacement of the flexible loop by TDC binding and formation of a hydrogen bond between TDC and S123, VtrA/VtrC is activated. However, when CDC binds to VtrA/VtrC, the protein complex is no longer primed for activation because the position of H50 is shifted upward by the presence of a coordinated water molecule between H50 and Y151 (**Figure 2A**).

However, the H50 pKa predictions do not provide an explanation for why VtrA/VtrC induces transcription upon binding TDC under pH 5.5 and pH 7.0 conditions, but not under pH 9.0 conditions (**Figure 5**). The predicted pKa of 3.7 for H50 in the TDC-bound structure is lower than the three pH conditions we tested. Based on this predicted pKa, H50 would be protonated in less than 50% of VtrA/VtrC dimers at pH 5.5. At pH 7.0 and pH 9.0, gradually lower proporitions of VtrA/VtrC would have protonated H50 residues. These values are more consistent with the pH titration of TDC binding measured by ITC for the periplasmic region. Of note, the structures used in the PROPKA predictions contained only the periplasmic domains of VtrA/VtrC. Since there are currently no structures of full-length VtrA/VtrC, the effect of different pH conditions on the whole protein complex is not clear.

As shown previously by Gotoh et al., conjugated forms of the secondary bile acid DC, such as TDC, are high inducers of *VtrB* transcription via VtrA/VtrC, while the unconjugated primary bile acids CDC and CA do not induce transcription (Gotoh et al., 2010). Meanwhile, conjugated primary bile acids and unconjugated DC induce intermediate levels of transcription. The high inducing bile acids have a 12α-OH and an amino acid (taurine or glycine) conjugated to the end of the carbon side chain (**Figure 1A**). The non-inducing bile acids have a 7α-OH instead of 12α-OH group and no conjugated amino acids on the carbon side chain. The intermediate-inducing bile acids have just one of either the 12α-OH or a conjugated carbon side chain. Therefore, the position of hydroxyl groups in the steroid nucleus and the presence of an amino acid conjugated to the carbon side chain are correlated with the ability of a bile acid to induce *VtrB* transcription via VtrA/VtrC.

As shown in **Figure 1**, the bile acids TDC and CDC can compete to induce or suppress *VtrB* transcription, respectively. Primary bile acids such as CDC are made in the liver and conjugated to taurine or glycine prior to secretion into the duodenum (Begley et al., 2005). Bacteria in the intestines can extensively modify these bile acids using reactions such as de-conjugation by bile salt hydrolases (BSHs), hydroxylation, or dehydroxylation to form unconjugated primary bile acids and secondary bile acids like DC (Begley et al., 2005). Bile acids are reabsorbed from the intestines and reconjugated in the liver (Begley et al., 2005, Hofmann, 1999), producing conjugated secondary bile acids like TDC. Modifications of bile acids by BSHs have various proposed benefits for gut bacteria, including providing a nutrient source through release of conjugated amino acids, strengthening the bacterial membrane through incorporation of bile acids and cholesterol, or reducing the acidic and detergent properties of bile acids that are toxic to bacteria (Begley et al., 2005, Foley, O’Flaherty et al., 2019). Recent studies have also shown the modification of bile acids by BSHs of different microbiome species can provide protection against colonization by enteric pathogens such as *V. cholerae* and *C. difficile* by altering the makeup of the bile acid pool in the intestines (Alavi, Mitchell et al., 2020, Foley, Walker et al., 2022). In the case of *V. parahaemolyticus*, higher levels of BSH activity in the intestines may produce a greater proportion of unconjugated primary bile acids like CDC that suppress T3SS2 expression. Therefore, we predict gut microbiota BSH activity and the makeup of the intestinal bile acid pool may also be important for determining whether *V. parahaemolyticus* passes through the host intestines as a friend or foe.

## Experimental Procedures

### Bacterial strains and cell culture

The *V. parahaemolyticus* POR1 *vtrB-3XFLAG* and POR1Δ*vtrC* strains were derived from POR1 (clinical isolate RIMD2210633 containing deletions of TDH toxins). The *vtrB-3XFLAG* strain contains an insertion of three tandem FLAG epitopes at the 3’ end of the native *vtrB* coding region, made using the Ori6K/SacB suicide vector pDM4 (Chimalapati, Lafrance et al., 2020). The POR1Δ*vtrC* strain contains a deletion of the *vtrC* coding sequence (nucleotides 34-486) (Li et al., 2016). Single amino acid substitution mutants of VtrC were generated using site directed mutagenesis of the vectors pACYC-Duet-VtrC/VtrA and pBAD-FLAG-*vtrC* (Li et al., 2016). Primers used for cloning are listed in **Table S3**. For vector induced expression of FLAG-*vtrC* variants under the control of the arabinose inducible promoter, pBAD-FLAG-*vtrC* plasmids were introduced into POR1Δ*vtrC* using standard triparental mating. Subsequently, a pRU1701(Karunakaran, Mauchline et al., 2005) derivative plasmid containing the 300 bp upstream of *vtrB* (pRU1701 *vtrB* −300bp) was introduced into the POR1Δ*vtrC* pBAD-FLAG-*vtrC* strains using *E. coli* S17-1 (λ*pir*). The *V. parahaemolyticus* POR1 *vtrB-3XFLAG* strain was cultured in Marine LB (MLB) broth (LB broth with 3% NaCl). Strains containing the pBAD and pRU1701 plasmids were cultured in MLB broth supplemented with 100 µg/mL gentamicin and 250 μg/mL kanamycin. *V. parahaemolyticus* cultures were grown at 30°C, or as otherwise indicated.

### Antibodies

FLAG antibody was purchased from Sigma-Aldrich (F3165).

### VtrB-3XFLAG expression assay

*V. parahaemolyticus* POR1 *vtrB-3XFLAG* strain was grown overnight in MLB medium at 30°C. The day of the experiment, new cultures were inoculated with OD_600_ = 0.3 in MLB using the overnight cultures and incubated at 30°C for 2.5 hours. Cultures were then induced with taurodeoxycholate (TDC) and chenodeoxycholate (CDC) at the indicated concentrations and incubated at 37°C for 1 hour. Bacterial cultures equivalent to OD_600_ = 1.0 were collected and cell pellets were resuspended in 2x protein sample buffer. Protein expression was detected by western blot analysis.

### GFP reporter assay

*V. parahaemolyticus* POR1Δ*vtrC* pBAD-FLAG-*vtrC* and pRU1701 strains were grown in MLB medium at 30°C overnight. The day of the experiment, new cultures were inoculated with OD_600_ = 0.6 in MLB supplemented with 1% arabinose using the overnight cultures and incubated at 30°C. After 2 hours of incubation, samples were prepared in triplicate by diluting the previous cultures to OD_600_ = 0.3 in MLB supplemented with 1% arabinose and either 100 μg/mL taurodeoxycholate (TDC) or 100 μg/mL chenodeoxycholate (CDC). Samples (200 μL) were transferred to a 96-well plate and incubated at 37°C in the plate reader. GFP fluorescence intensity (FI) was measured every 5 minutes using the BMG Labtech CLARIOstar^Plus^ plate reader with an excitation filter of 470 nm and emission filter of 515 nm. Absorbance at 600 nm (A_600_) was measured after each GFP FI measurement and these values were used to normalized GFP FI to cell density. Plates were shaken in between each GFP FI and A_600_ measurement.

### Protein expression and purification

The periplasmic domain of VtrC (aa 31-161) and mutants with single amino acid mutations were co-expressed with the periplasmic domain of VtrA (aa 161-253) using variants of the pACYC-Duet-VtrC/VtrA construct in which VtrC has an N-terminal hexahistidine tag(Li et al., 2016). The constructs were expressed in *E. coli* BL21(DE3) cells and purified using nickel-affinity and size exclusion chromatography (SEC)(Li et al., 2016). Briefly, all cultures were grown in LB at 37°C until OD_600_ = 0.5 – 0.6 and induced with 0.4 mM isopropyl β-D-thiogalactopyranoside (IPTG) overnight at 16°C. Cells were lysed in buffer A (50 mM Tris pH 8.0 and 100 mM NaCl) supplemented with 1 mM phenylmethylsulfonyl fluoride (Sigma) using a cell disrupter (Emulsiflex C3, Avestin Inc.). Clarified and filtered (0.45 μm pore size) lysates were incubated with Qiagen Ni-NTA resin (30210) for 30 minutes at 4°C with nutation. Lysate and resin were applied to a gravity column. Protein-bound resin was washed and eluted with buffer A supplemented with 15 mM (wash) and 250 mM (elution) imidazole, as described previously (Li et al., 2016). Eluted proteins were further purified by SEC on a Superdex 200 Increase 10/300 GL column (Cytiva) with buffer A. For crystallographic studies, the VtrA/VtrC heterodimer bound to the bile acid CDC was purified by nickel-affinity chromatography with 0.5 mM CDC in all buffers. The final SEC purification was performed with buffer B (10 mM Tris pH 8.0 and 10 mM NaCl) supplemented with 0.5 mM CDC. CDC-bound protein was buffer exchanged into buffer B without bile acid prior to crystallographic studies.

### Crystallization and X-ray data collection

Crystals of the VtrA/VtrC periplasmic domain heterodimer bound to the bile acid CDC were grown using the hanging-drop vapor diffusion method from drops containing 1 μL of protein (4.8 mg/mL) and 1 μL of reservoir solution (16% polyethylene glycol (PEG) 3,350, 0.2 M calcium chloride) and equilibrated over 250 μL of reservoir solution. Crystals typically appeared in 2-7 days at 20°C and grew to their maximal extent by 1 week. Crystals were cryoprotected by transferring to a final solution of 18% PEG 3,350, 0.2 M calcium chloride, 10 mM Tris pH 8.0 and 35% glycerol, and flash-cooled in liquid nitrogen.

Data were collected at APS beamline 19-ID at 100 K, and were indexed, integrated, and scaled using the HKL-3000 program package (Minor, Cymborowski et al., 2006). CDC-bound VtrA/VtrC crystals belonged to space group C2 with unit cell parameters of a = 142.60 Å, b = 41.76 Å, c = 168.96 Å and β = 91.57° and contained four molecules each of VtrA/VtrC heterodimer per asymmetric unit, with a solvent content of 45%. Data collection statistics are provided in Table 2.

**Table 2.**
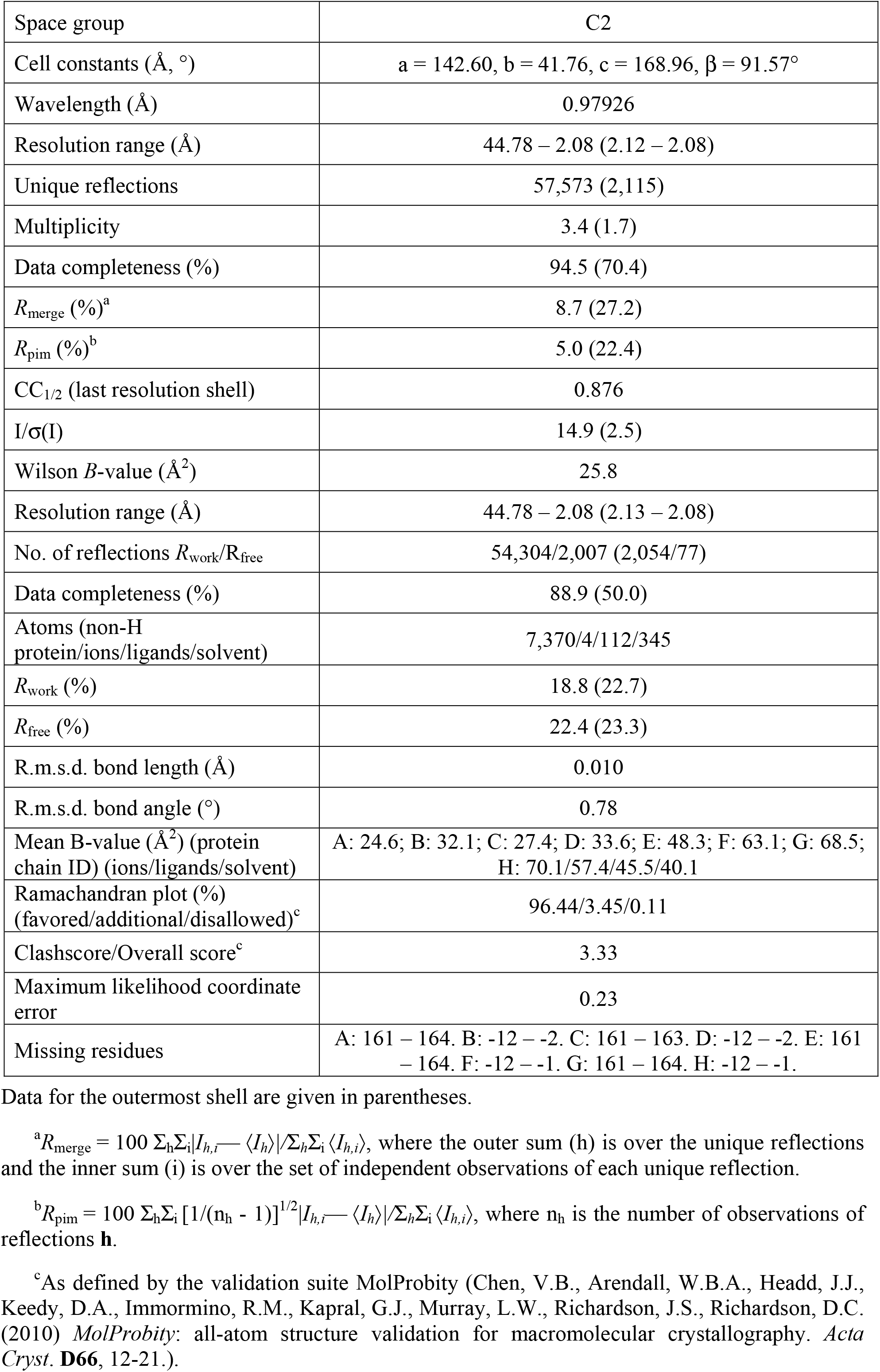
Data collection and refinement statistics, VtrA/C and CDC complex

### Phase determination and structure refinement

Phases for the CDC-bound VtrA/VtrC heterodimer were obtained by the molecular replacement method in the program *Phaser* (McCoy, Grosse-Kunstleve et al., 2007) using the coordinates for the TDC-bound VtrA/VtrC heterodimer (5KEW). Manual model building was performed in the program *Coot* (Emsley, Lohkamp et al., 2010). Positional and isotropic atomic displacement parameter (ADP) and TLS ADP refinement was performed to a resolution of 2.08 Å using the program *Phenix* (Afonine, Mustyakimov et al., 2010) with a random 3.70% of all data set aside for an R_free_ calculation. The current model contains four VtrA/VtrC heterodimers in the asymmetric unit, each bound to one molecule of CDC, as well as four calcium ions and 292 water molecules. A *Molprobity* (Chen, Arendall et al., 2010) generated Ramachandran plot indicates 96.4% of residues are in the most favored regions and 0.1% (one residue) is in disallowed regions. Phasing and model refinement statistics are provided in Table 2.

### Isothermal titration calorimetry (ITC)

All variants of the VtrA/VtrC periplasmic domain complex were dialyzed against the assay buffer (50 mM Tris pH 8.0, 100 mM NaCl) overnight at 4°C. Taurodeoxycholic acid (TDC) and chenodeoxycholic acid (CDC) were each prepared by dissolving dry powder (Sigma) with the same dialysis buffer to a concentration of 200 μM. ITC experiments were performed at 25°C on a MicroCal iTC200 system (Malvern), with reference power at 5 μcal/s and stirring rate at 750 rpm. Measurements were generally performed as 21 injections of 200 μM TDC or CDC (1 μL for the first injection and 2 μL for injections 2-21) into approximately 200 μL of 20 μM VtrA/C. ITC data were integrated and analyzed using NITPIC 1.3.0 (Keller, Vargas et al., 2012, Scheuermann & Brautigam, 2015) and SEDPHAT version 15.2b (Brautigam, Zhao et al., 2016). ITC data plots were prepared with GUSSI 1.4.2 (Brautigam, 2015).

For ITC experiments performed under pH 7.0 and pH 9.0 conditions, the VtrA/VtrC periplasmic domain complex was purified as described above, except the final purification by SEC was performed using pH 7.0 (50 mM Tris pH 7.0, 100 mM NaCl) and pH 9.0 (50 mM Tris pH 9.0, 100 mM NaCl) buffers. Protein and bile acid samples for ITC were prepared as above using the pH 7.0 and pH 9.0 buffers.

## Data Availability

Structure factors and coordinates for the VtrA/VtrC and CDC complex were deposited in the Protein Data Bank (PDB) under the accession code 8DML.

## Supporting Information

This article contains supporting information (Figures S1, S2, S3, S4 Tables S1, S2, S3).

## Acknowledgments

We thank the Orth Lab for valuable discussion and advice, Dr. Chad Brautigam and Dr. Shih-Chia Tso for assistance with ITC data collection and analysis, and the Structural Biology Lab for support with X-ray crystallographic studies. The structure in this report is derived from work performed on beamline 19-ID at the Argonne National Laboratory, Structural Biology Center at the Advanced Photon Source, operated by UChicago Argonne, LLC, for the US Department of Energy, Office of Biological and Environmental Research under contract DE-AC02-06CH11357.

## Funding and Additional Information

This work was funded by the Welch Foundation grant I-1561 (K.O.), Once Upon a Time…Foundation (K.O.), National Institutes of Health Grant T32GM131963 (A.Z.) and R01 GM115188 (K.O.). The content is solely the responsibility of the authors and does not necessarily represent the official views of the National Institutes of Health.

## Supplemental Figures

**Figure S1.**
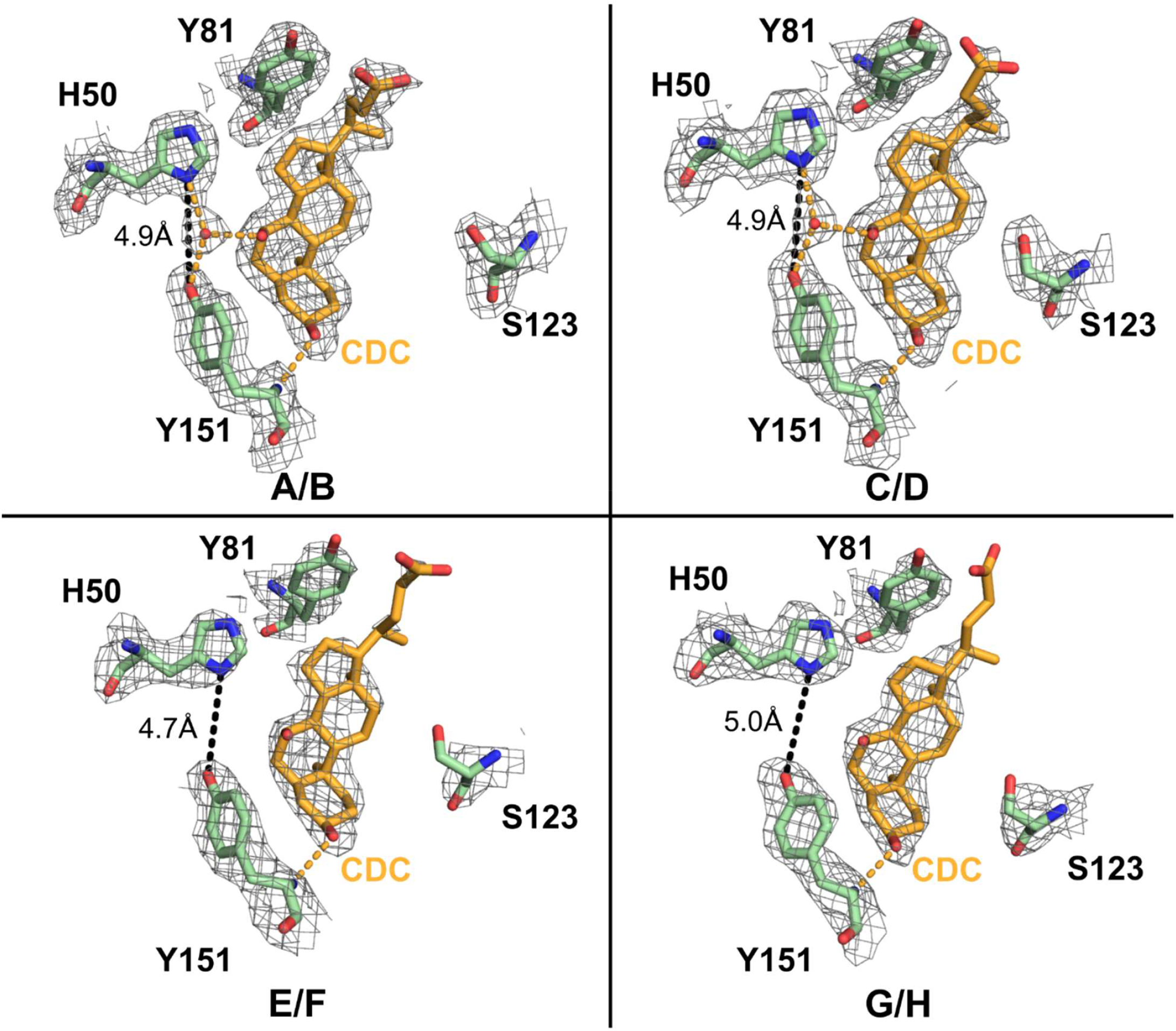
Electron density around CDC molecules and binding pocket residues. Kicked F_o_-F_c_ omit maps of the four CDC molecules bound to the A/B, C/D, E/F, and G/H heterodimers. CDC molecules (orange) and binding pocket residues (light green) are shown as sticks. Modeled water molecules are shown as red spheres. Hydrogen bonds are shown as orange dashed lines. Distance measurements are indicated with black dashed lines. Maps are shown as grey mesh (contoured at the 1σ level) and carved around sticks at a 1.6 Å radius.

**Figure S2.**
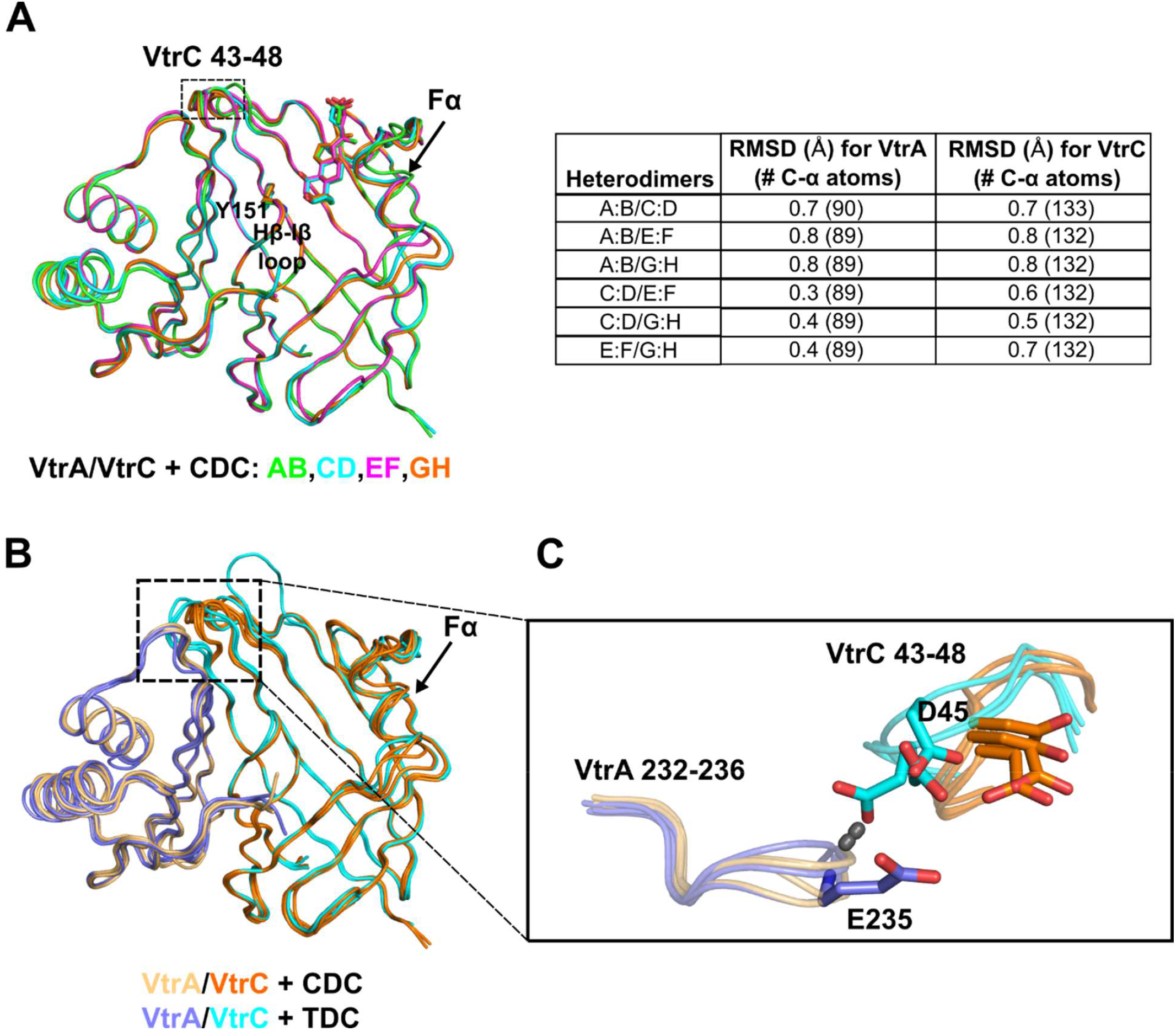
Alignment of the CDC-bound structure heterodimers and comparison with TDC-bound structure heterodimers. **(A)** (Left) Superposition of ribbon models for all four heterodimers in the asymmetric unit of the CDC-bound crystal. VtrC Y151 side chain and CDC shown as sticks. (Right) Root-mean-square-deviations (RMSD) for each superposition in (Å) with the number of aligned α-carbon atoms in parentheses. Superpositions were made via the DaliLite server (http://ekhidna.biocenter.helsinki.fi/dali_lite/start) (Holm et al., 2000). **(B)** Superposition of (A) with the three heterodimers in the asymmetric unit of the TDC-bound crystal as ribbons. **(C)** “Top” view of VtrA/VtrC interface boxed in (B). VtrA/VtrC modelled as ribbons. VtrA E235 and VtrC D45 side chains shown as sticks. Hydrogen bond between the backbone amide of VtrA E235 and the side chain of VtrC D45 shown as a dashed cyan line.

**Figure S3.**
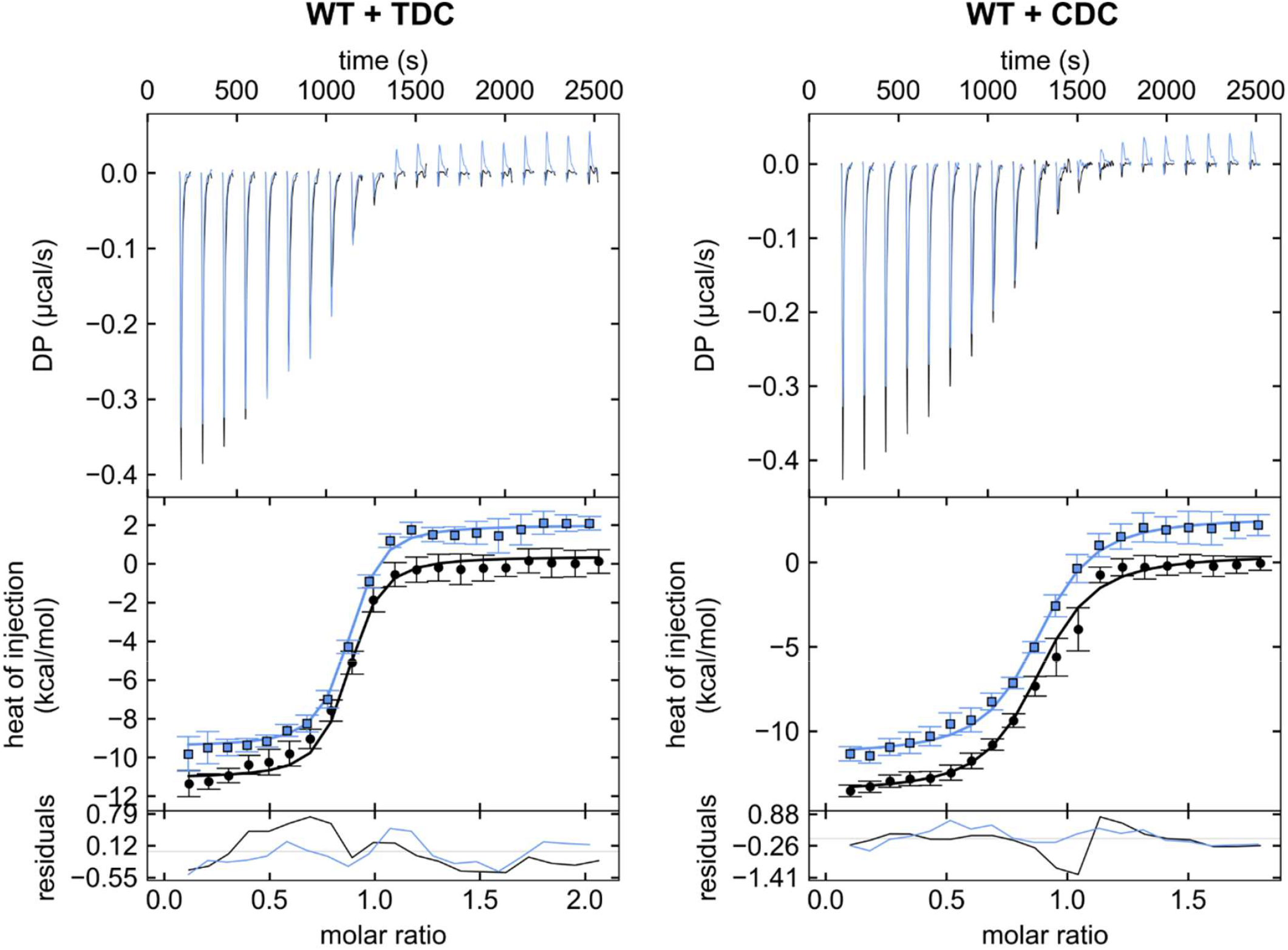
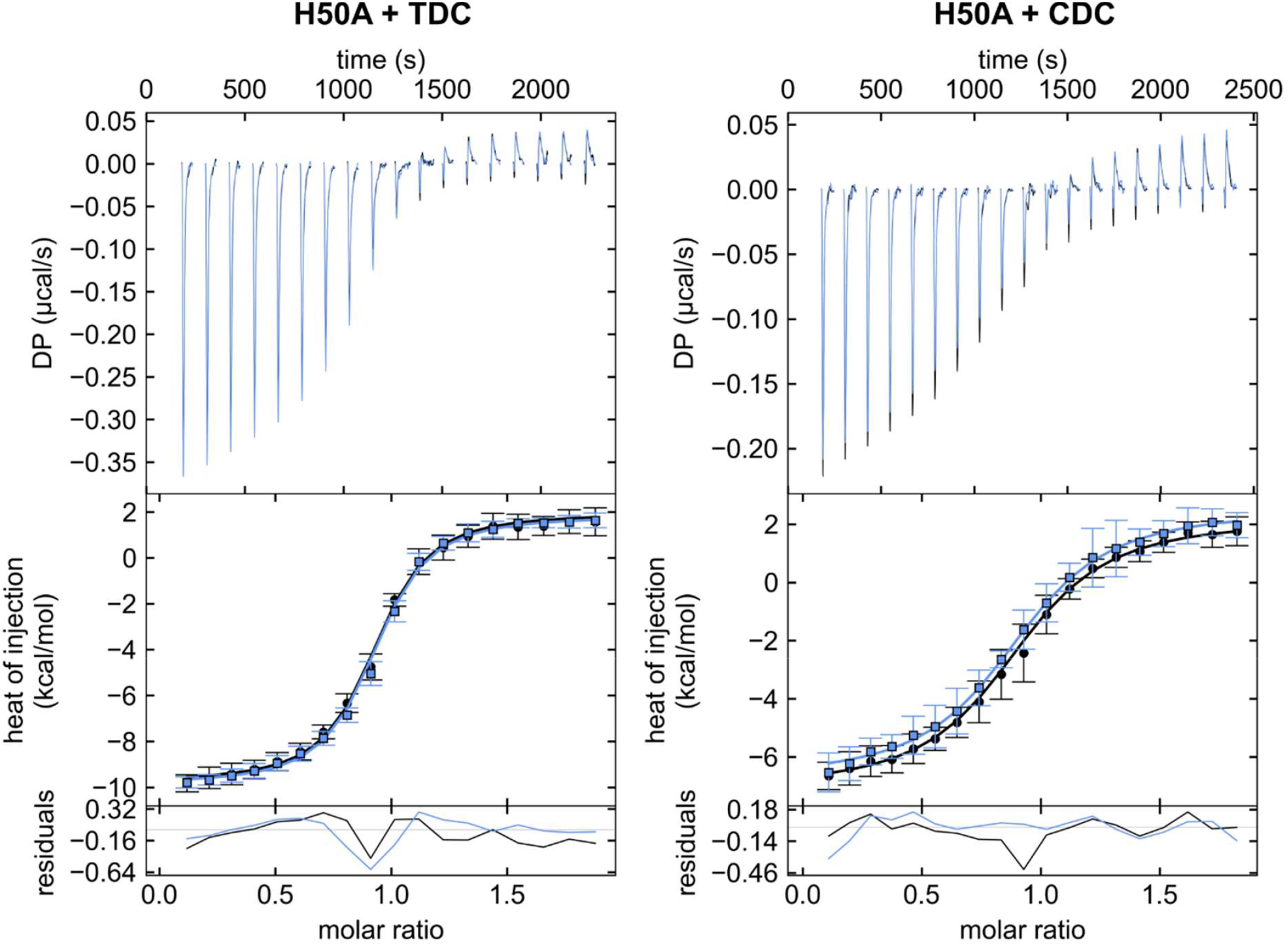
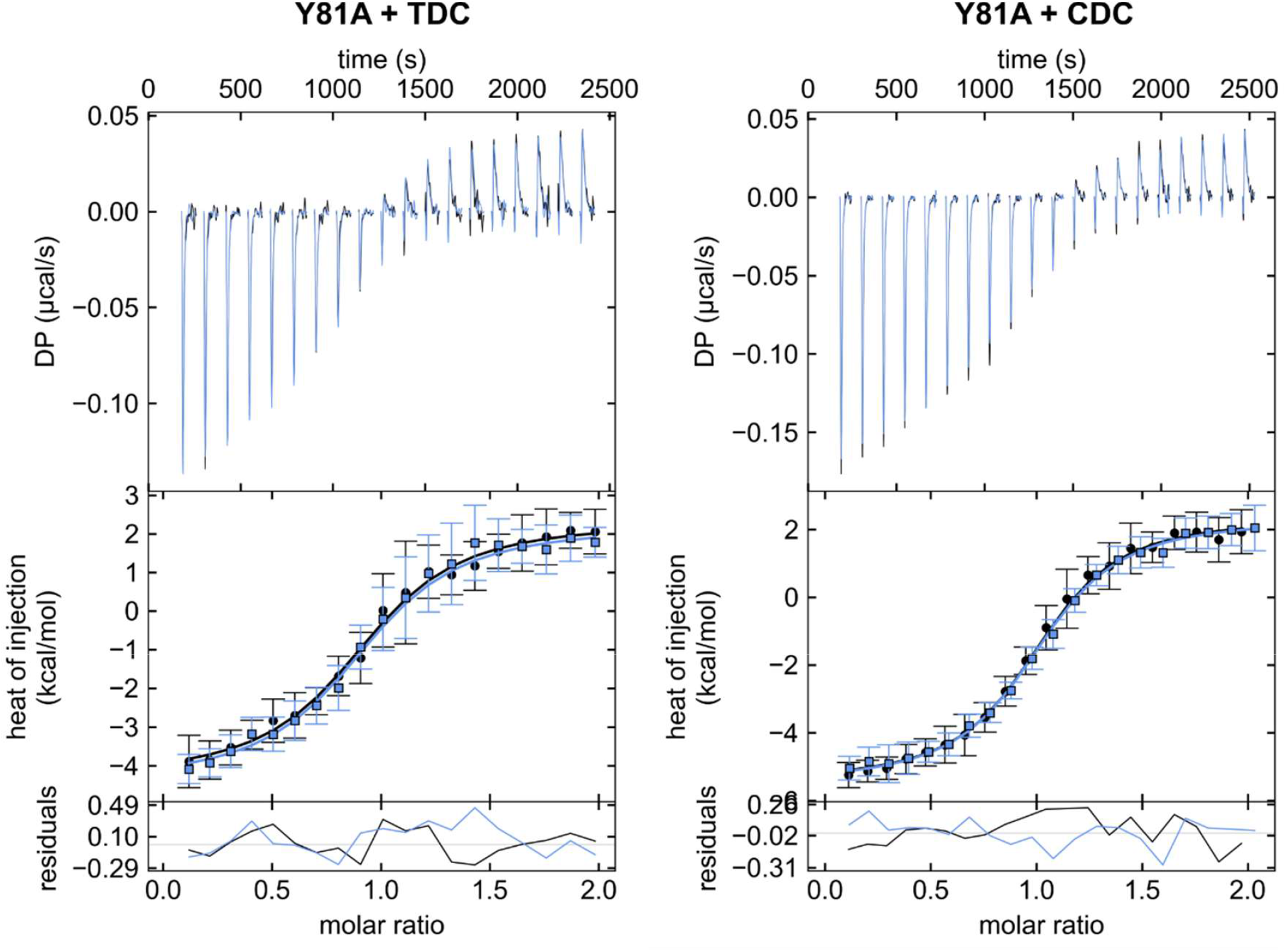
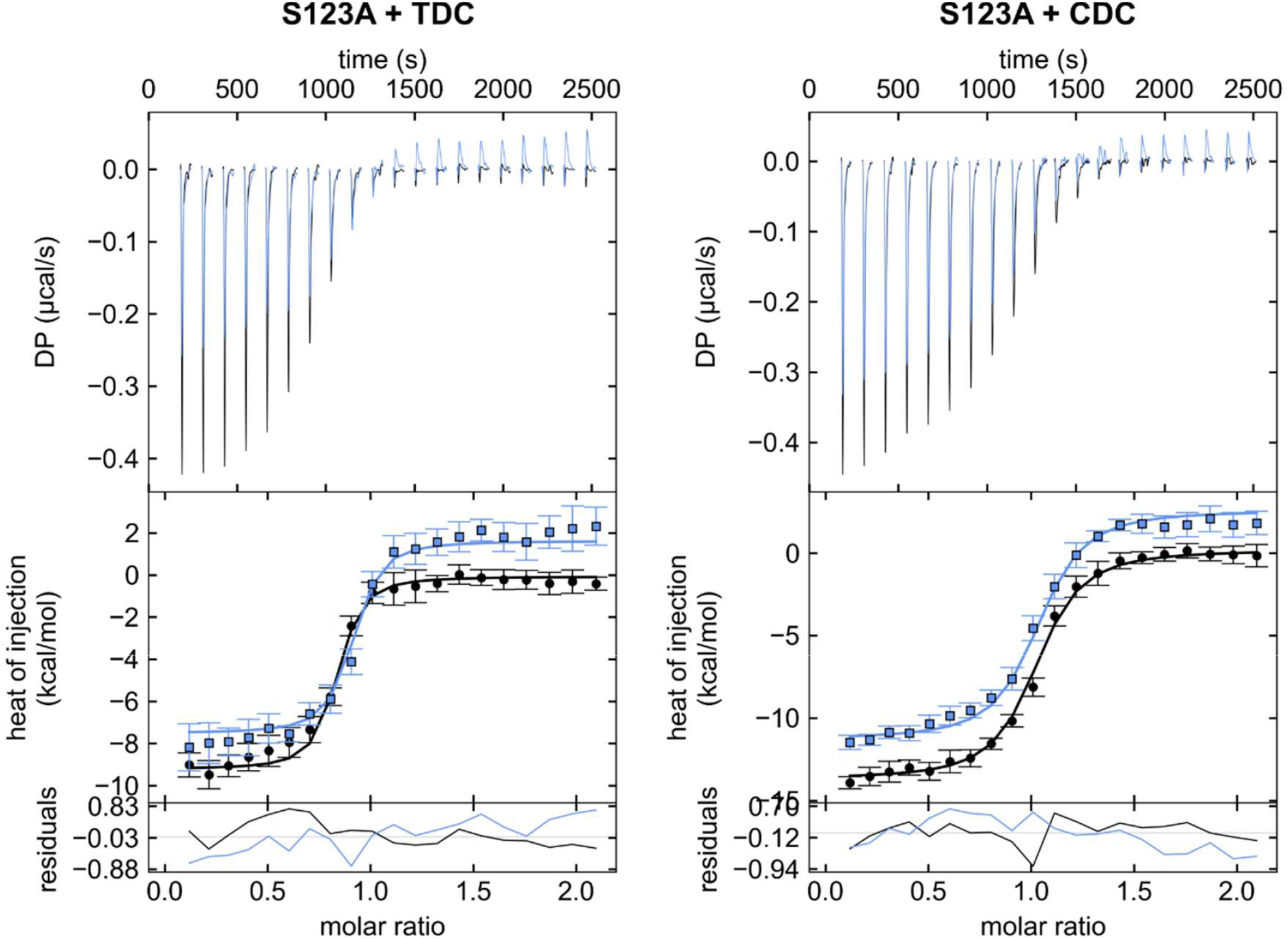
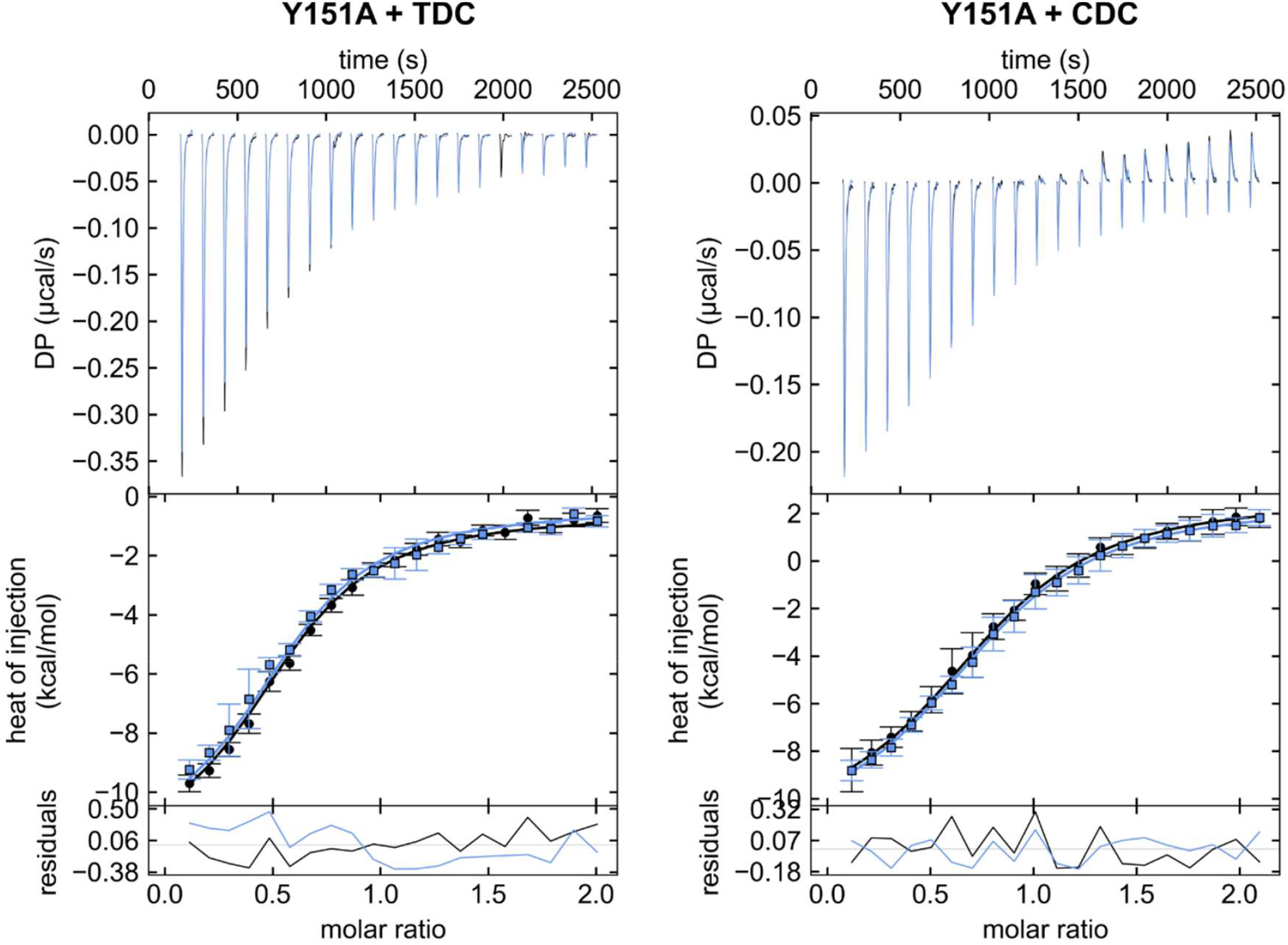
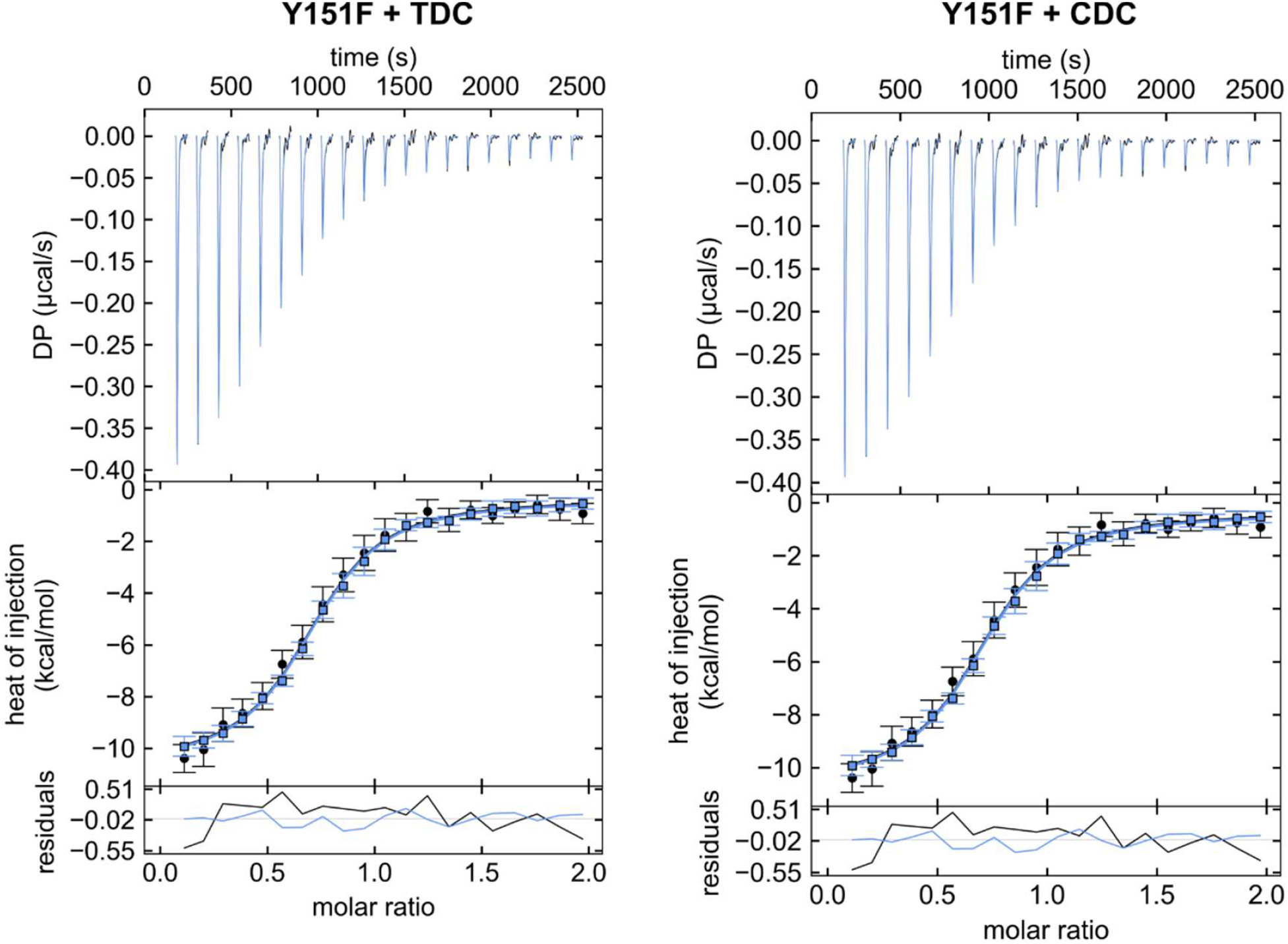
ITC thermograms for binding of TDC and CDC to VtrA/VtrC constructs. Thermodynamic parameters (Table 1, Table S1) were determined by global fitting of duplicate isotherms (presented in black and blue).

**Figure S4.**
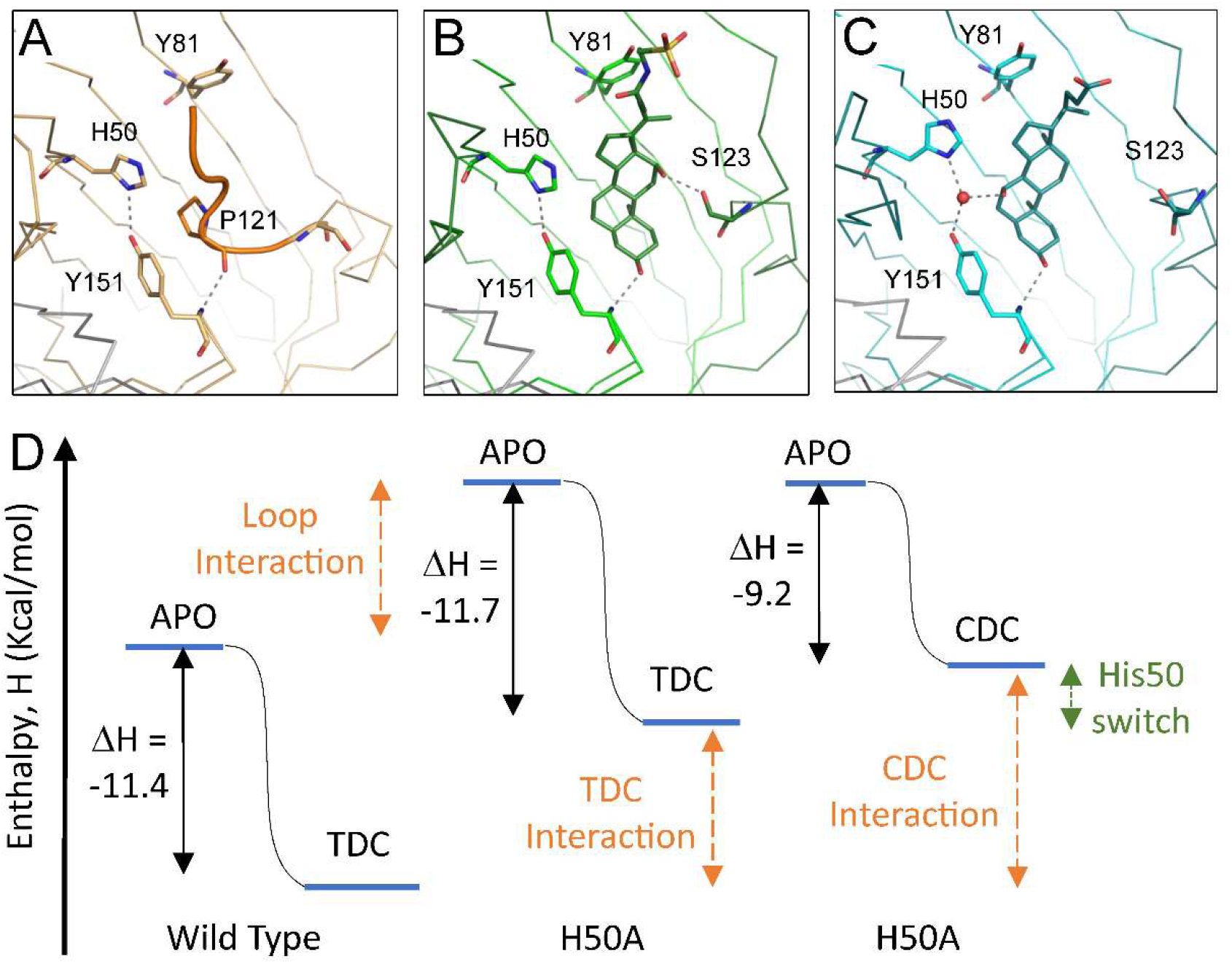
Entropic contribution to specificity binding switch. **A)** Apo VtrA/VtrC structure (orange ribbon), with labeled key binding pocket residues (stick). Hydrogen bonds are indicated by gray dotted lines. The bile acid binding pocket is covered by the Fα extended loop (in orange tube), with the ring of P121 replacing the middle 6-member ring of the bile acid steroid backbone and the P121 backbone OH replacing the bile acid 3α-OH. H50 and Y181 form a similar hydrogen bond as seen in **B)** the structure (green ribbon) bound to TDC (dark green stick). The 12α-OH of TDC forms a hydrogen bond with Ser123 in the Fα helix (dark green ribbon). **C)** H50 conformation switch in structure (cyan ribbon) bound to CDC (teal stick) establishes a hydrogen bonding network to the 7α-OH of CDC through an ordered water (red sphere). **D** Enthalpy coordinates for wild type and H50A (labeled below) binding. Reaction from apo (left bar) to bound (TDC/CDC labeled bars) has a similar change for wild type and H50A binding to TDC (left black arrows), suggesting a similar shift of both reactants and products to higher enthalpic energy for the mutation and a similar enthalpic energy for binding the Fα extended loop (orange arrow) and the TDC (orange arrow). The change in enthalpy for CDC binding to wild type is smaller, resulting in an increased enthalpic energy (green arrow) that might reflect the conformation switch.

**Table S1.**
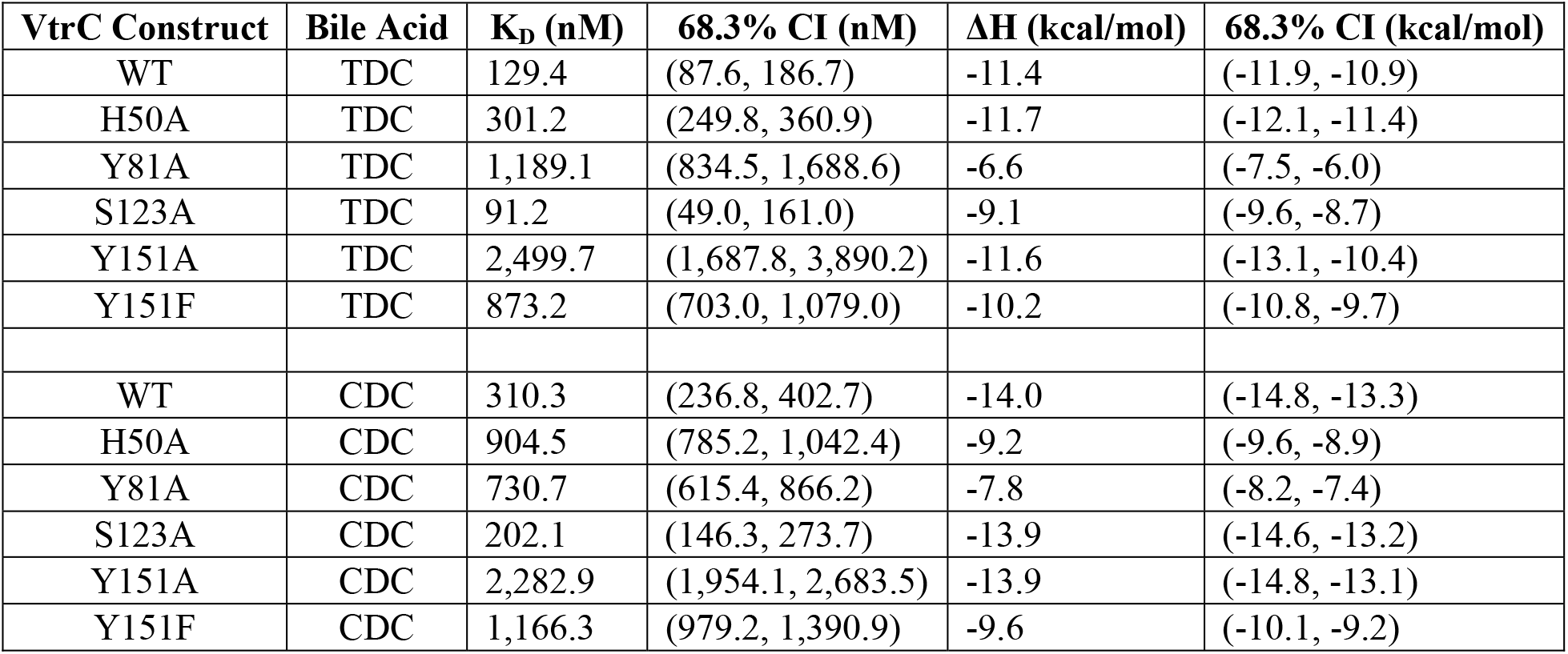
Thermodynamic parameters with 68.3% confidence intervals (CI) of TDC and CDC binding to various VtrA/VtrC constructs measured by ITC.

**Table S2.**
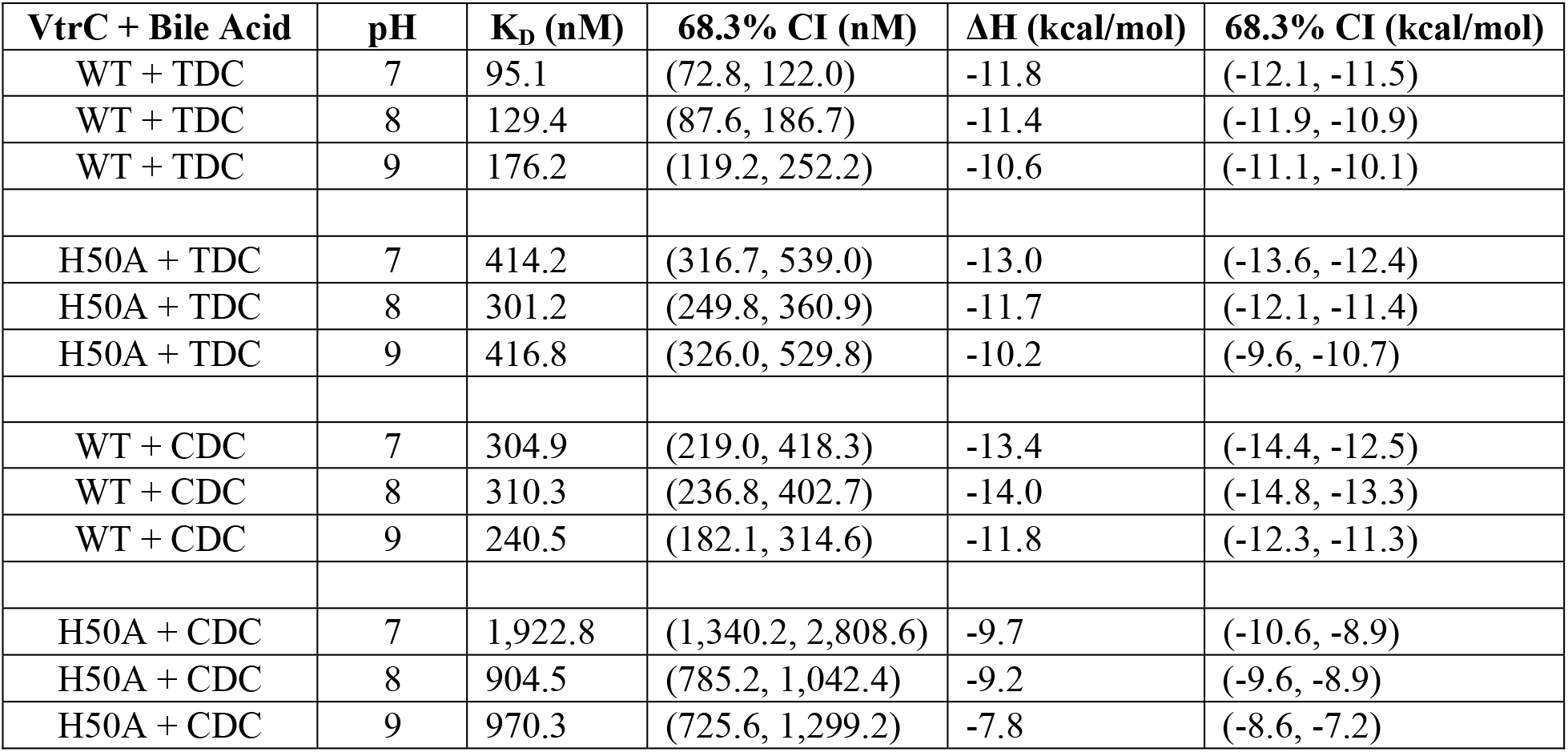
Thermodynamic parameters with 68.3% confidence intervals (CI) of TDC and CDC binding to VtrA/VtrC constructs under different pH conditions measured by ITC.

**Table S3.**
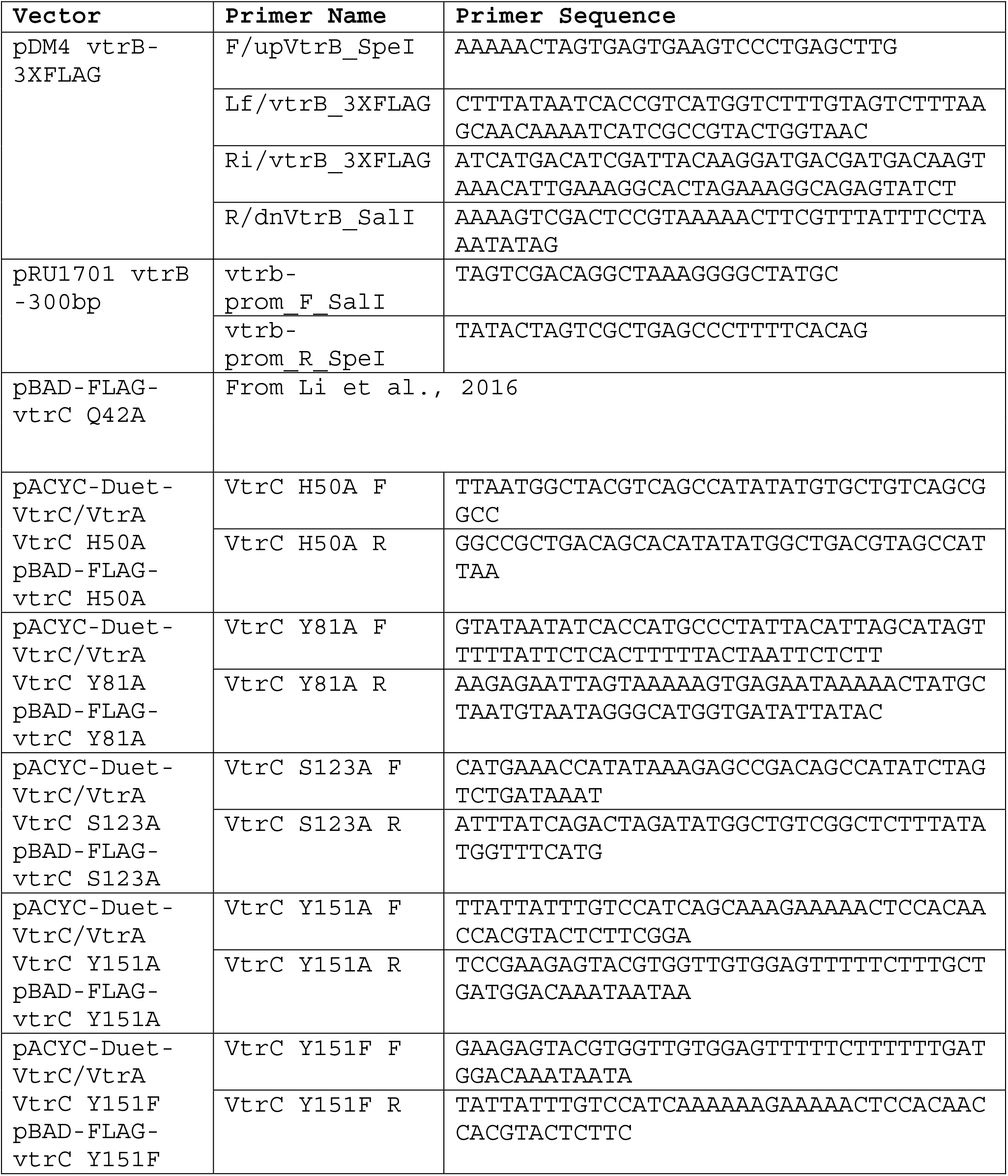
Primers used in this study.

## References

Afonine PV, Mustyakimov M, Grosse-Kunstleve RW, Moriarty NW, Langan P, Adams PD (2010) Joint X-ray and neutron refinement with phenix.refine. Acta Crystallogr D Biol Crystallogr 66: 1153–63

Alavi S, Mitchell JD, Cho JY, Liu R, Macbeth JC, Hsiao A (2020) Interpersonal Gut Microbiome Variation Drives Susceptibility and Resistance to Cholera Infection. Cell 181: 1533-1546.e13

Begley M, Gahan CG, Hill C (2005) The interaction between bacteria and bile. FEMS Microbiol Rev 29: 625–51

Brautigam CA (2015) Chapter Five - Calculations and Publication-Quality Illustrations for Analytical Ultracentrifugation Data. In Methods in Enzymology, Cole JL (ed) pp 109-133. Academic Press

Brautigam CA, Zhao H, Vargas C, Keller S, Schuck P (2016) Integration and global analysis of isothermal titration calorimetry data for studying macromolecular interactions. Nature Protocols 11: 882–894

Chen VB, Arendall WB, 3rd, Headd JJ, Keedy DA, Immormino RM, Kapral GJ, Murray LW, Richardson JS, Richardson DC (2010) MolProbity: all-atom structure validation for macromolecular crystallography. Acta Crystallogr D Biol Crystallogr 66: 12–21

Chimalapati S, Lafrance AE, Chen L, Orth K (2020) Vibrio parahaemolyticus: Basic Techniques for Growth, Genetic Manipulation, and Analysis of Virulence Factors. Curr Protoc Microbiol 59: e131

de Souza Santos M, Orth K (2014) Intracellular Vibrio parahaemolyticus escapes the vacuole and establishes a replicative niche in the cytosol of epithelial cells. mBio 5: e01506–14

de Souza Santos M, Salomon D, Li P, Krachler A-M, Orth K (2015) 8 - Vibrio parahaemolyticus virulence determinants. In The Comprehensive Sourcebook of Bacterial Protein Toxins (Fourth Edition), Alouf J, Ladant D, Popoff MR (eds) pp 230–260. Boston: Academic Press

Emsley P, Lohkamp B, Scott WG, Cowtan K (2010) Features and development of Coot. Acta Crystallogr D Biol Crystallogr 66: 486–501

Evans DF, Pye G, Bramley R, Clark AG, Dyson TJ, Hardcastle JD (1988) Measurement of gastrointestinal pH profiles in normal ambulant human subjects. Gut 29: 1035–41

Foley MH, O’Flaherty S, Barrangou R, Theriot CM (2019) Bile salt hydrolases: Gatekeepers of bile acid metabolism and host-microbiome crosstalk in the gastrointestinal tract. PLoS Pathog 15: e1007581

Foley MH, Walker ME, Stewart AK, O’Flaherty S, Gentry EC, Allen G, Patel S, Pan M, Beaty VV, Vanhoy ME, Dougherty MK, McGill SK, Gulati A, Dorrestein PC, Baker ES, Redinbo MR, Barrangou R, Theriot CM. (2022) Distinct bile salt hydrolase substrate preferences dictate C. difficile pathogenesis. bioRxiv. 10.1101/2022.03.24.485529 [preprint]

Gotoh K, Kodama T, Hiyoshi H, Izutsu K, Park K-S, Dryselius R, Akeda Y, Honda T, Iida T (2010) Bile Acid-Induced Virulence Gene Expression of Vibrio parahaemolyticus Reveals a Novel Therapeutic Potential for Bile Acid Sequestrants. PLOS ONE 5: e13365

Hofmann AF (1999) The Continuing Importance of Bile Acids in Liver and Intestinal Disease. Archives of Internal Medicine 159: 2647–2658

Jacob-Dubuisson F, Mechaly A, Betton JM, Antoine R (2018) Structural insights into the signalling mechanisms of two-component systems. Nat Rev Microbiol 16: 585–593

Karunakaran R, Mauchline TH, Hosie AHF, Poole PS (2005) A family of promoter probe vectors incorporating autofluorescent and chromogenic reporter proteins for studying gene expression in Gram-negative bacteria. Microbiology (Reading) 151: 3249–3256

Keller S, Vargas C, Zhao H, Piszczek G, Brautigam CA, Schuck P (2012) High-Precision Isothermal Titration Calorimetry with Automated Peak-Shape Analysis. Analytical Chemistry 84: 5066–5073

Kinch LN, Cong Q, Jaishankar J, Orth K (2022) Co-component signal transduction systems: Fast-evolving virulence regulation cassettes discovered in enteric bacteria. Proc Natl Acad Sci U S A 119: e2203176119

Kodama T, Gotoh K, Hiyoshi H, Morita M, Izutsu K, Akeda Y, Park K-S, Cantarelli VV, Dryselius R, Iida T, Honda T (2010) Two Regulators of Vibrio parahaemolyticus Play Important Roles in Enterotoxicity by Controlling the Expression of Genes in the Vp-PAI Region. PLOS ONE 5: e8678

Li H, Robertson AD, Jensen JH (2005) Very fast empirical prediction and rationalization of protein pKa values. Proteins: Structure, Function, and Bioinformatics 61: 704–721

Li P, Rivera-Cancel G, Kinch LN, Salomon D, Tomchick DR, Grishin NV, Orth K (2016) Bile salt receptor complex activates a pathogenic type III secretion system. Elife 5: e15718

McCoy AJ, Grosse-Kunstleve RW, Adams PD, Winn MD, Storoni LC, Read RJ (2007) Phaser crystallographic software. J Appl Crystallogr 40: 658–674

Minor W, Cymborowski M, Otwinowski Z, Chruszcz M (2006) HKL-3000: the integration of data reduction and structure solution--from diffraction images to an initial model in minutes. Acta Crystallogr D Biol Crystallogr 62: 859–66

Okada R, Matsuda S, Iida T (2017) Vibrio parahaemolyticus VtrA is a membrane-bound regulator and is activated via oligomerization. PLOS ONE 12: e0187846

Olsson MHM, Søndergaard CR, Rostkowski M, Jensen JH (2011) PROPKA3: Consistent Treatment of Internal and Surface Residues in Empirical pKa Predictions. Journal of Chemical Theory and Computation 7: 525–537

Scheuermann TH, Brautigam CA (2015) High-precision, automated integration of multiple isothermal titration calorimetric thermograms: New features of NITPIC. Methods 76: 87–98

Schlundt A, Buchner S, Janowski R, Heydenreich T, Heermann R, Lassak J, Geerlof A, Stehle R, Niessing D, Jung K, Sattler M (2017) Structure-function analysis of the DNA-binding domain of a transmembrane transcriptional activator. Sci Rep 7: 1051

Søndergaard CR, Olsson MH, Rostkowski M, Jensen JH (2011) Improved Treatment of Ligands and Coupling Effects in Empirical Calculation and Rationalization of pKa Values. J Chem Theory Comput 7: 2284–95

Zhang L, Krachler AM, Broberg CA, Li Y, Mirzaei H, Gilpin CJ, Orth K (2012) Type III effector VopC mediates invasion for Vibrio species. Cell Rep 1: 453–60

Holm L, Park J (2000) DaliLite workbench for protein structure comparison. Bioinformatics 16:566–567

